# Highly biased agonism for GPCR ligands via nanobody tethering

**DOI:** 10.1101/2023.10.10.561766

**Authors:** Shivani Sachdev, Brendan A. Creemer, Thomas J. Gardella, Ross W. Cheloha

## Abstract

Ligand-induced activation of G protein-coupled receptors (GPCRs) can initiate signaling through multiple distinct pathways with differing biological and physiological outcomes. There is intense interest in understanding how variation in GPCR ligand structure can be used to promote pathway selective signaling (“biased agonism”) with the goal of promoting desirable responses and avoiding deleterious side effects. Here we present a new approach in which a conventional peptide ligand for the type 1 parathyroid hormone receptor (PTHR1) is converted from an agonist which induces signaling through all relevant pathways to a compound that is highly selective for a single pathway. This is achieved not through variation in the core structure of the agonist, but rather by linking it to a nanobody tethering agent that binds with high affinity to a separate site on the receptor not involved in signal transduction. The resulting conjugate represents the most biased agonist of PTHR1 reported to date. This approach holds promise for facile generation of pathway selective ligands for other GPCRs.

## Introduction

G protein-coupled receptors (GPCRs) are the largest family of cell surface proteins and the most common target of approved therapeutics. These receptors are activated by diverse molecules and stimuli ranging from proteins and peptides to small molecules, photons, and protons^1^. Recent structural studies using X-ray crystallography and cryo-electron microscopy have provided unprecedented molecular level insights into the structural changes associated with receptor activation^2,3^. A handful of common signatures have been observed in structures of activated receptors in complex with G proteins or ligands relative to unactivated counterparts^4^. Even among the receptor structures classified as “active state”, subtle structural differences are observed among receptors bound to different ligands, G-proteins, and accessory proteins^5^.

A wide body of experimental findings has demonstrated that different compounds that bind to and activate the same receptor induce divergent biological and physiological responses in a paradigm known as ligand bias or pathway selective signaling^6,7^. Biased ligands induce preferential signaling through one receptor transducer (such as Gαs-mediated adenylate cyclase activation) over another (such as β-arrestin) relative to an index comparator ligand^8^. Extensive effort has been invested to link the subtle differences in receptor structure induced by the binding of different ligands to biased signaling outputs^9^. In some examples there is a clear connection between variation in the receptor conformation induced by the binding of a biased ligand and the type of signaling bias observed, though this is not true for all examples^10,11^. As such, prospective efforts for the rational design of biased agonists with desired signaling bias profiles are challenging.

The GPCR superfamily is divided into separate classes based on sequence conservation and the presence of class-specific structural features^12^. Class B1 GPCRs are characterized by a large extracellular domain (ECD), which facilitates the binding of medium-sized polypeptide ligands^2^. The interaction between class B1 GPCRs and their ligands has traditionally been described as a two-site mode of binding^13^. In this two-site mechanism, the interaction between the C-terminal portion of the peptide ligand and the ECD of the receptor (site 1 interaction) provides most of the binding affinity and specificity for the bimolecular interaction. Site 1 also serves to anchor the N-terminal portion of the peptide in a position where it can contact the transmembrane portion of the receptor (site 2 interaction, orthosteric site), induce a conformational change, and initiate a signaling response. Although the site 2 interaction is low affinity, it is hypothesized to be singularly responsible for ligand agonist behavior, including the induction of pathway selective signaling^14–18^. In the two-site model, the ECD-peptide C-terminus interaction serves as a tethering point to improve ligand affinity and potency but plays no direct role in inducing a conformational change. By extension, structural modifications in the ligand C-terminus are predicted to substantially impact ligand binding but not pathway selective signaling, although recent findings suggest this is an oversimplification^19,20^.

The interaction of the type-1 parathyroid hormone receptor (PTHR1), a class B1 GPCR, with its ligands has served as an exemplar of this two-site mechanism of ligand binding. PTHR1 is bound by two naturally occurring peptide hormones, parathyroid hormone (PTH) and PTH-related protein (PTHrP), with full biological activity residing in N-terminal 34 or 36 residues respectively^21^. Activation of PTHR1 induces signaling through Gα proteins (primarily Gαs and secondarily Gαq) and β-arrestin. Modifications in the C-terminal portion of PTH_1-34_ have strong effects on receptor binding and signaling potency (site 1) whereas N-terminal modifications mostly impact receptor activation efficacy and pathway selectivity (site 2). For example, modifications at positions 23, 24, and 28 in PTH_1-34_ diminish binding to the receptor ECD and signaling potency without a substantial impact on ligand efficacy or pathway selectivity^16,22,23^. In contrast, modifications within the first 11 residues of PTH have been identified that affect ligand agonist efficacy, pathway signaling selectivity, and signaling localization^16–18,24,25^.

Some optimized PTH_1-11_ analogues exhibit signaling profiles and pathway selectivity similar to PTH_1-34_, albeit with somewhat reduced potency and affinity^24,26^. Past work showed that such truncated PTH analogues could be linked to single domain antibodies (nanobodies, Nbs) using a combination of enzymatic protein labeling, solid phase peptide synthesis, and chemoselective conjugation chemistry^27^. This novel approach termed “CLAMP”, uses Nbs, which are the smallest antibody fragments that maintain the desirable characteristics of conventional antibodies such as high target affinity and specificity^28^. In contrast to conventional antibodies, Nbs can be produced in high yield from bacteria, exhibit high stability even in the absence of glycosylation and disulfide bond formation, and are comprised of a single polypeptide chain, which facilitates straightforward construction of multi-specific conjugates. Past work has shown that the linkage of a PTH_1-11_ analogue with a Nb that bound to PTHR1 extracellular domain augmented signaling through the Gαs pathway^27^. The binding of Nb to ECD can be envisioned as a surrogate for the site 1 interaction engaged by the PTH_1-34_ C-terminal region. The conventional two-site model would suggest that PTH_1-11_-Nb conjugates would exhibit agonist properties (signaling efficacy, pathway selectivity, and signaling localization) similar to PTH_1-11_ alone.

Here we test this hypothesis by synthesizing a variety of Nb-PTH_1-11_ conjugates through chemical, enzymatic, and recombinant protein expression methods. Using a panel of Nbs, including a newly characterized PTHR1-binding Nb (Nb_PTHR1-X2_), we show that PTH_1-11_-Nb conjugates are unexpectedly highly selective for Gαs/cAMP pathway activation. This finding is in stark contrast to conventional PTHR1 ligands, such as PTH_1-11_ and PTH_1-34_, which signal through all PTHR1-engaged pathways. Mechanistic studies revealed that the PTH_1-11_-Nb conjugates can activate signaling through a mode that involves two receptor protomers (termed “activation *in trans”*), which is qualitatively distinct from that of conventional ligands of PTHR1. These findings show that agonist properties, such as pathway selectivity, for class B1 GPCRs are dependent not only on the structure of the ligand that engages the orthosteric site (site 2) but also the positioning of the high affinity tethering interaction (site 1). This finding holds important implications for efforts to develop potent biased agonists.

## Results

We sought to probe the consequences of outsourcing the receptor binding function of PTHR1 ligands to artificial building blocks and non-natural binding sites. Towards this end we synthesized a set of ligands designed to target either WT PTHR1 or engineered receptors (Figure 1A). Nb-ligand conjugates were prepared using recombinant Nb expression, site-specific labeling, peptide synthesis, and click chemistry (Figure 1B). All peptides, including PTH_1-11_-6E, which contains a six-carbon linker (6-aminohexanoic acid, Ahx) between components, were synthesized using standard Fmoc-based solid phase peptide synthesis. Peptide and conjugate identity were confirmed by mass spectrometry (Supporting Tables 1 and 2). Previously described variants of PTHR1 (Figure 1C) engineered to contain a high affinity binding site for an epitope tag-binding Nb (Nb_6E_ binds PTHR1-6E) or an epitope peptide (6E peptide binds PTHR1-Nb_6E_) were also deployed^27,29^. Either epitope tag (6E) or Nb (Nb_6E_) were engrafted into a flexible and unstructured region of the receptor encoded by exon 2, which is known to be dispensable for high affinity ligand binding^30^. Through this design, the signaling properties of PTH_1-34_ could be compared to engineered ligands. For example, both PTH_1-34_ and PTH_1-11_-6E are predicted to engage in a high affinity interaction with PTHR1-Nb_6E_ but with different mechanisms (Figure 1C). PTH_1-34_ affinity comes mostly through engagement of a cleft in the receptor extracellular domain by the PTH_12-34_ fragment, whereas PTH_1-11_-6E instead binds to the engrafted Nb_6E_. Analogously, both Nb_6E_-PTH_1-11_ and PTH_1-34_ were predicted to exhibit high affinity binding at PTHR1-6E.

**Figure 1:**
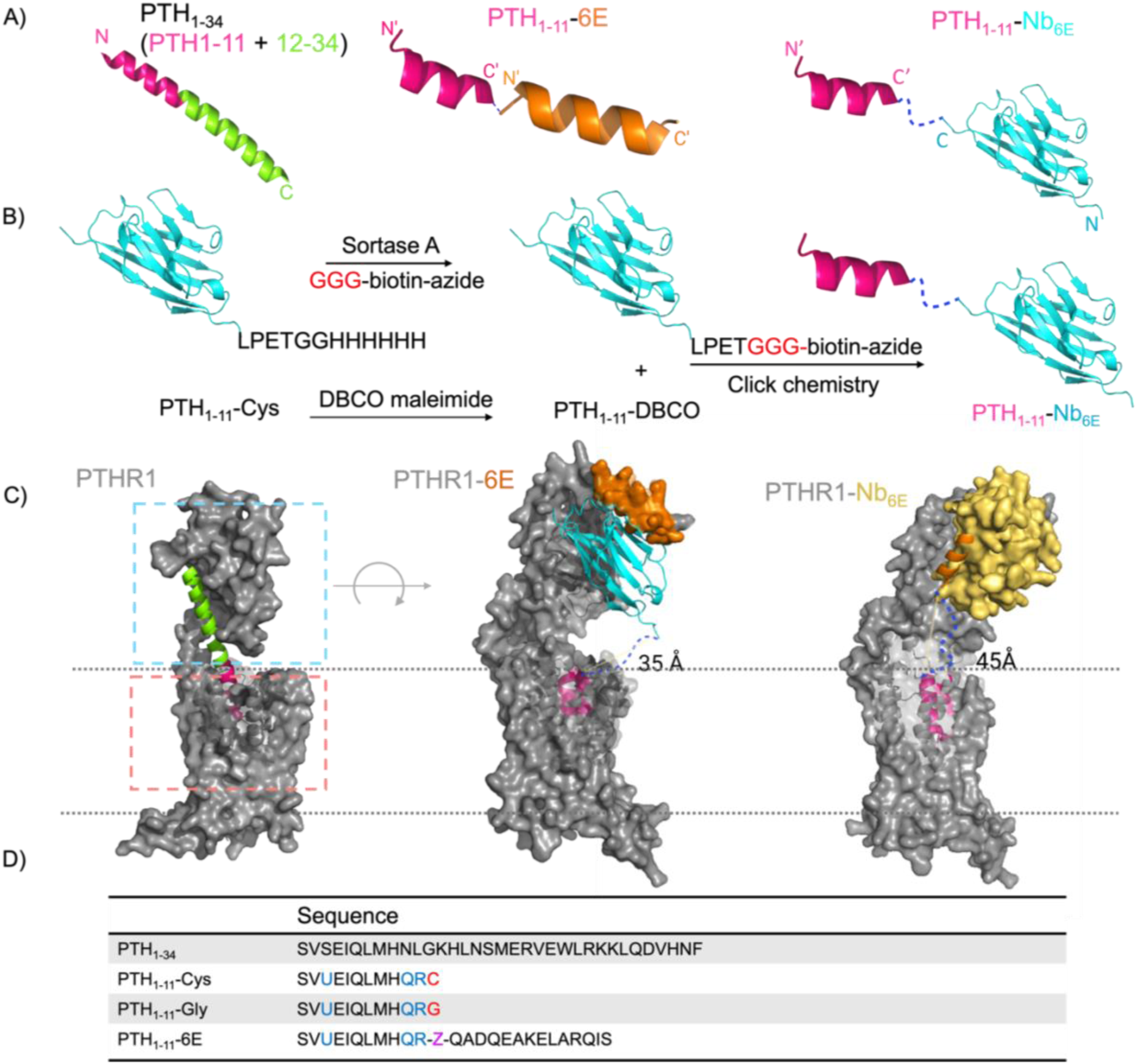
Ligand-nanobody conjugate synthesis and receptor designs. A) Schematic representation of the ligands used in this study. B) Synthetic scheme for preparation of Nb-ligand conjugates. C) Variation on the canonical two-site binding model of Class B GPCR activation. For PTHR1, the model is based on the published crystal structure of PTH_1-34_ bound to PTHR1 (PDB: 6FJ3). The peptide agonist-ECD interaction is highlighted in the blue dashed box (site 1) and the agonist-transmembrane domain interaction is shown in salmon dashed box (site 2). Models of PTHR1-6E and PTHR1-Nb_6E_ interacting with binding partners were generated using Alphafold2 (see Methods) and are shown rotated relative to wild type PTHR1. PTHR1-6E is shown bound to PTH_1-11_-Nb_6E_ conjugate (pink and cyan). PTHR1-Nb_6E_ is shown bound PTH_1-11_-6E (pink and orange). The dark blue dashed lines depict a hypothetical linker between the C-terminus of PTH_1-11_ and Nb_6E_ or 6E in these models. Distances listed were measured in Alphafold generated models using Pymol. D) Sequences of ligands used in this study. Sites with mutations relative to natural PTH_1-34_ are highlighted with colored text. The residue abbreviated with “U” corresponds aminoisobutyric acid; “Z” corresponds to 6-aminohexanoic acid. Mass spectrometry characterization of compounds is shown in Supporting Tables 1 and 2. Unless stated otherwise, reference to “PTH_1-11_” in the remainder of the text refers to PTH_1-11_-Gly, whereas Nb-PTH_1-11_ conjugates are constructed using PTH_1-11_-Cys.

**Table 1.** Pharmacological parameters for ligands and conjugates across all the functional assays tested. Values shown correspond to mean (SEM) from the number of replicates indicated in Supporting Table 3. Dashes (“-“) indicate that the EC_50_ value could not be determined due to weak ligand activity in that assay. “ND” indicates that the assay was not conducted.

| EC<sub>50</sub> (±SEM)<br>Ligands | cAMP | Washout<br>AUC | Gαs | β-arrestin 2<br>(plasma<br>membrane) | β-arrestin 2<br>(endosome) |
| --- | --- | --- | --- | --- | --- |
| **PTHR1-6E** |  |  |  |  |  |
| PTH<sub>1-34</sub> | 25 (8) | 36 (8) | 0.3 (0.1) | 189 (48) | 142 (62) |
| PTH<sub>1-11</sub> | 50 (23) | 125 (57) | 32 (13) | 11300 (3100) | 4800 (900) |
| PTH<sub>1-11</sub>-Nb<sub>6E</sub> | 22 (7) | 8 (0.7) | 5 (0.7) | - | - |
| PTH<sub>1-11</sub>-Nb<sub>neg</sub> | - | - | - | - | - |
| **PTHR1-Nb<sub>6E</sub>** |  |  |  |  |  |
| PTH<sub>1-34</sub> | 0.4 (±0.1) | 1.2 (0.4) | 1.1 (0.8) | 106 (72) | 54 (19) |
| PTH<sub>1-11</sub> | 28 (11) | 240 (72) | 198 (142) | 27200 (7200) | 23100 (1400) |
| PTH<sub>1-11</sub>-Ahx-6E | 0.07 (0.05) | 0.2 (0.08) | 4.4 (3) | - | - |
| **PTHR1** |  |  |  |  |  |
| PTH<sub>1-34</sub> | 5 (2) | 5.4 (1.7) | 0.1 (0.06) | 62 (12) | 25 (3) |
| PTH<sub>1-11</sub> | 22 (5) | 342 (116) | 26 (2.6) | 12300 (1700) | 11900 (4200) |
| PTH<sub>1-11</sub>-Nb<sub>PTHR1</sub> | 3 (0.4) | 4 (0.6) | 2.3 (0.9) | - | - |
| PTH<sub>1-11</sub>-Nb<sub>PTHR1X2</sub> | 5.3 (1.5) | 2.3 (1) | ND | - | ND |

Experiments were performed to assess how signaling characteristics varied in response to alterations in the mode of interaction between the receptor and ligand conjugates. PTHR1 signals primarily through the Gαs pathway, which induces intracellular cyclic adenosine monophosphate (cAMP) production. Previously developed cell lines, derived from HEK293, which stably express individual PTHR1 variants of interest and a luciferase-based biosensor used to monitor cyclic adenosine monophosphate (cAMP) production were applied^27,31^ . We compared the activities of PTH_1-34_, PTH_1-11_, and PTH_1-11_-conjugates in cell lines expressing either PTHR1-6E or PTHR1-Nb_6E_. For both cell lines PTH_1-34_ exhibited a typical PTHR1 cAMP dose-response pattern (Fig. 2), with EC_50_ values in the low nM range (Table 1). As expected, the PTH_1-11_ peptide fragment, which lacks residues important for ECD binding, showed a reduced potency compared to full-length PTH_1-34_. Conjugation of PTH_1-11_ with Nb_6E,_ which binds to the 6E epitope in the ECD of PTHR1-6E, caused a substantial enhancement in potency in cells expressing this receptor (Fig. 2, Table 1), in agreement with past findings^27^. In contrast, the conjugation of PTH_1-11_ to a negative control nanobody (PTH_1-11_-Nb_Neg_), which recognizes an epitope not present (BC2)^32^, provided conjugates with no activity at PTHR1-6E (Fig. 2). We also evaluated the persistence of cAMP generation for conjugates following removal of free ligand (“washout”). Past work has shown that ligands that exhibit prolonged PTHR1 washout responses also induce prolonged physiological activity *in vivo*^20^. Our data reveal that PTH_1-11_-Nb_6E_ induced cAMP responses that were similar in magnitude and duration to PTH_1-34_, remaining stable throughout the course of 30 minutes (Fig. 2). Signaling duration was also evaluated through measurement of the area under the curve (AUC) after washout, also in agreement with past findings^27^. By this measure, PTH1-11-Nb_6E_ induced more enduring washout responses than PTH_1-34_, whereas responses from PTH_1-11_ were weaker (Fig. 2, Table 1).

**Figure 2:**
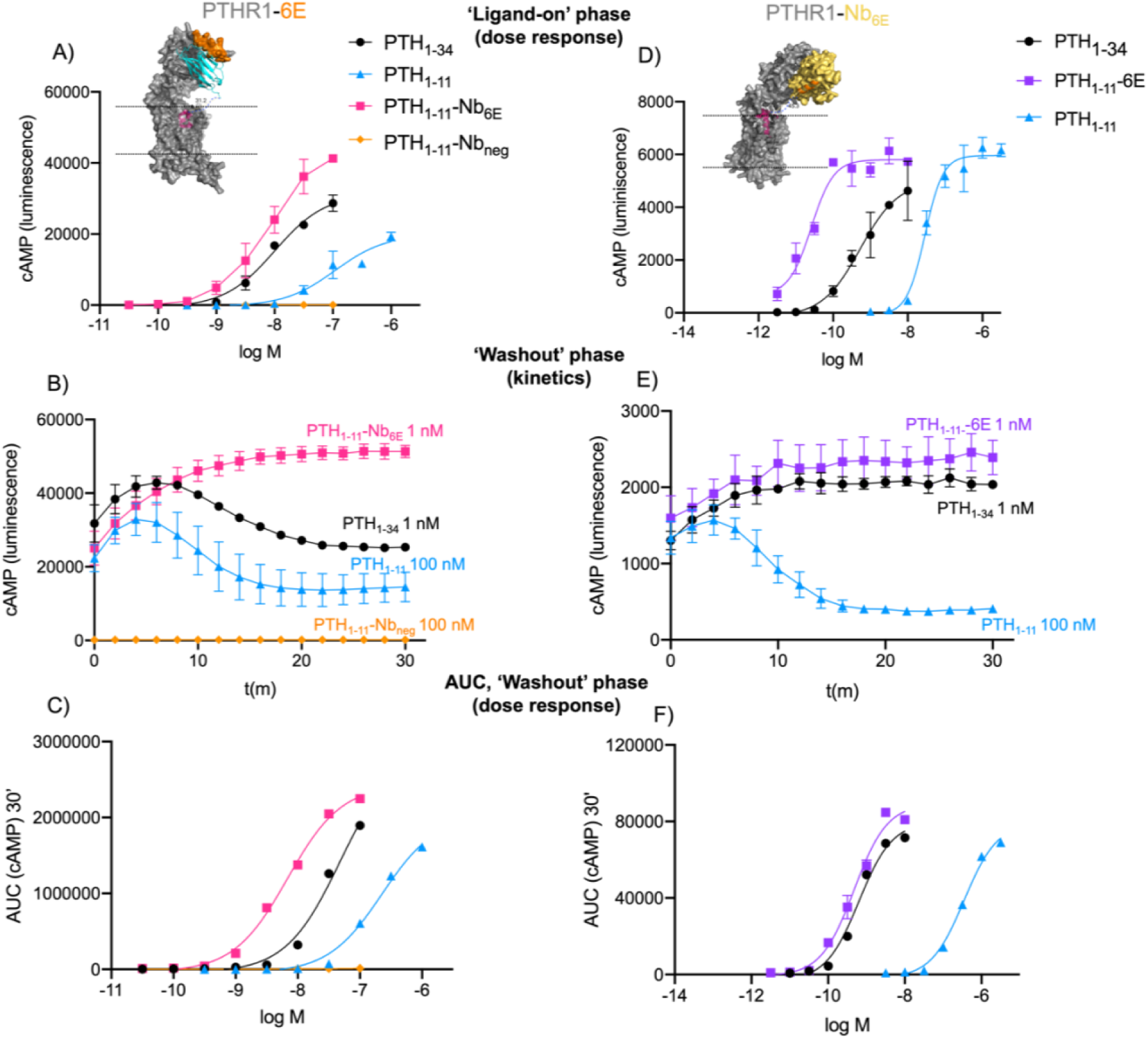
Ligand-induced cAMP responses in engineered receptors. Data correspond to experiments run using cells expressing PTHR1-6E (panels A-C) or PTHR1-Nb_6E_ (panels D-F). A) Concentration-response curves for cAMP production in cells expressing PTHR1-6E. Consistent color coding for ligands is used throughout the figure. B) Representative kinetic plots for ligand-induced signaling following the removal of the unbound ligand in a washout assay. C) Quantitation of duration of washout responses summarized as AUC. Panels D-F show analogous data for cells expressing PTHR1-Nb_6E._ Data points correspond to mean and associated SD from technical replicates. Curves were generated using a three-parameter logistic sigmoidal model. Tabulation of agonist potency parameters are shown in Table 1, which are derived from 3-5 independent experiments.

Similar experiments were performed on the cell line expressing PTHR1-Nb_6E_. PTH_1-11_-6E more effectively induced cAMP production than PTH_1-11_, with a 400-fold difference in potency (Table 1). In washout assays, PTH_1-11_-6E induced more enduring cAMP responses relative to the other peptides tested. To assess whether the prolonged signaling from PTH_1-11_-6E corresponded with continuous engagement of receptor, or whether two site binding permitted repeated association/dissociation cycles at different receptor sites, we evaluated the impact of antagonists added at the beginning of the washout phase (Supporting Figure 1). PTH_1-11_-6E washout responses were highly sensitive to inhibitors. Addition of synthetic 6E peptide causes inhibition through competition with PTH_1-11_-6E for Nb_6E_ binding, whereas SW106 interferes with the binding of PTH ^33^. The marked effects of both types of inhibitors on PTH -6E signaling suggests that this ligand does not continuously engage both of its binding sites throughout washout. The synthetic 6E peptide competitor is cell impermeable; its effective antagonism in this context suggests much of the signaling of PTH_1-11_-6E originates from the cell surface. Similarly high levels inhibitor sensitivity were previously observed for analogues of PTH_1-34_ that specifically signal from the cell surface^18,25^.

To support these findings, we conducted a distinct assay, based on the use of a bioluminescence resonance energy transfer (BRET) biosensor^34^, to measure ligand-induced Gαs dissociation from the plasma membrane. This experimental format does not rely on the signal amplification inherent in the cAMP-based assay described above. In this assay, PTH_1-11_-Nb_6E_ or PTH_1-11_-6E exhibited potency and efficacy comparable to PTH_1-34_ on PTHR1-6E or PTHR1-Nb_6E_, respectively (Fig. 3A), in line with their behavior in the cAMP assay.

**Figure 3.**
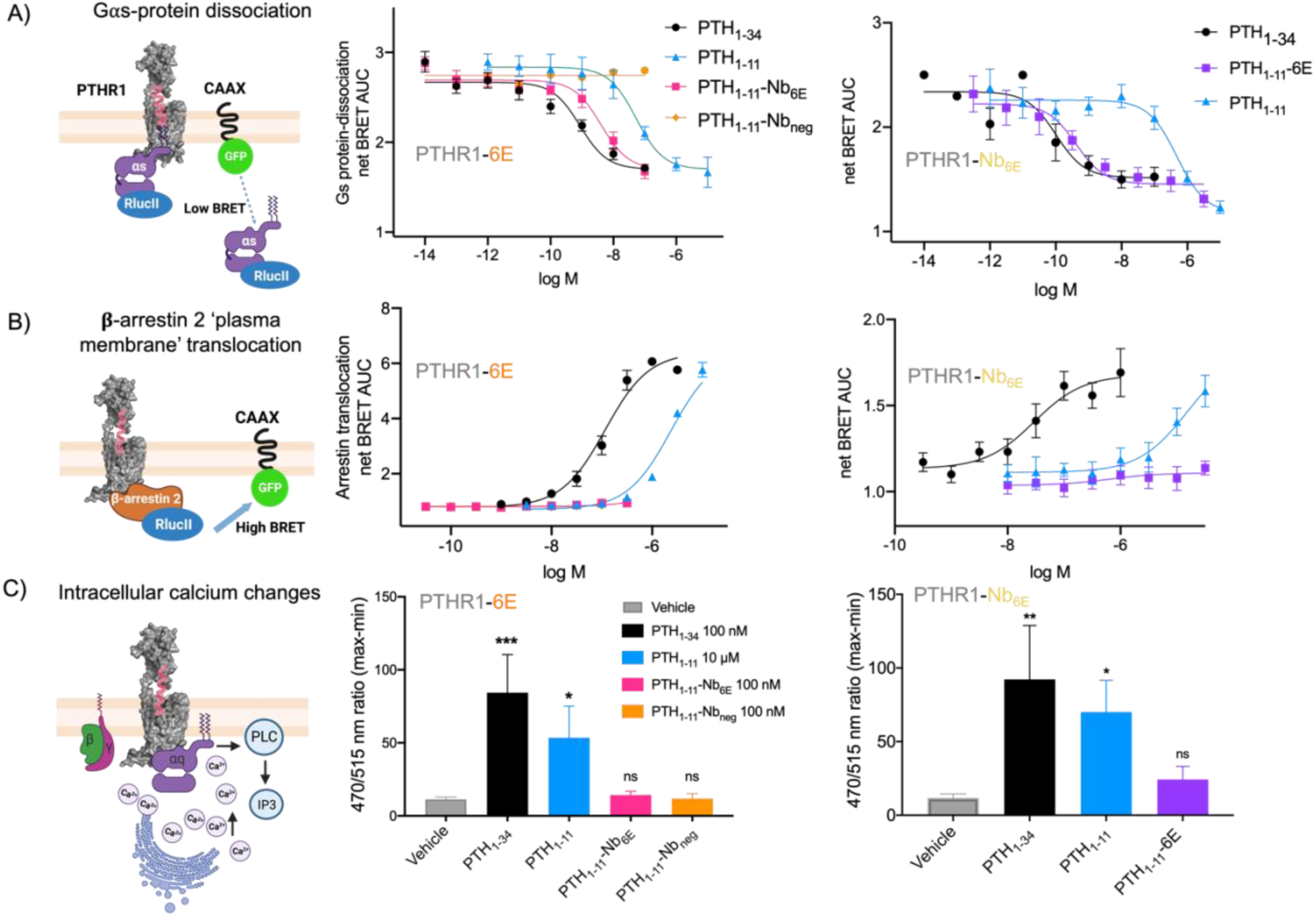
Ligand-induced signaling in engineered receptors through diverse pathways. A) Schematic and data corresponding to a BRET assay used to monitor Gαs-activation. A decrease in BRET ratio corresponds to ligand-induced dissociation of heterotrimeric Gαs protein complex from the plasma membrane. B) Schematic and data corresponding to a BRET assay used to monitor β-arrestin 2 recruitment. An increase in BRET ratio indicates ligand-induced arrestin translocation to the plasma membrane. In panels A-B, the data correspond to AUC values generated from kinetic measurements. Data points correspond to mean ± SD from technical replicates in a representative experiment. Characterization of endosomal β-arrestin 2 recruitment is shown in Supporting Figure 2. C) Schematic and data corresponding to the assay used to measure calcium mobilization induced by engagement of the Gαq-PTHR1 pathway. Cells were incubated with the Calbryte 520 AM Ca^2+^ indicator dye and exposed to indicated saturating concentrations of ligands. Data are presented as the relative fluorescence intensity normalized to signal background as means ± SEM from 3-5 independent experiments conducted with six technical replicates per experiment. Statistical significance was assessed by one-way ANOVA, with Dunnett’s post hoc correction (\**p* < 0.05; \*\**p* < 0.01; \*\*\**p* < 0.001; \*\*\*\**p* < 0.0001; ns not significant). Compiled and quantified data from this figure are shown in Table 1.

We also measured ligand-induced recruitment of β-arrestin 2 to the plasma membrane and to early endosomes using a bystander BRET assay^34^. In each assay, PTH_1-34_ showed the expected dose-response for induction of β-arrestin 2 recruitment (Fig. 3B-C, Table 1), with somewhat diminished potency compared to cAMP assays. PTH_1-11_ was less potent relative to PTH_1-34_. Strikingly, PTH_1-11_-Nb_6E_ displayed a drastically impaired capacity to induce recruitment of β-arrestin 2 to the plasma membrane or to early endosomes in cells stably expressing PTHR1-6E (Fig. 3, Supporting Figure 2). Assays in cells expressing PTHR1-Nb_6E_ showed that PTH_1-11_-6E was nearly inactive for inducing β-arrestin 2 recruitment, even at a concentration as high as 30 µM (Fig. 3B-C). A comparable lack of activity was recorded for PTH_1-11_-Nb_6E_ and PTH_1-11_-6E for inducing Gαq-mediated calcium mobilization in relevant cell lines (Figure 3C). The capacity of PTH_1-11_-Nb_6E_ and PTH_1-11_-6E to robustly induce cAMP responses in relevant cell lines, without concomitant induction of β-arrestin 2 recruitment or Gαq activation, corresponds to a clearcut example of highly biased agonism.

This finding prompted us to investigate whether the biased agonism profile observed with engineered receptors would translate to the native human PTHR1 receptor (hPTHR1). This required the use of Nbs that bind directly to hPTHR1. Sequences for Nbs that recognize PTHR1 (named here Nb_PTHR1_ and Nb_PTHR1-**X2**_) have been reported^35^, albeit with sparse characterization. Nb_PTHR1_ was previously used (with the name VHH_PTHR_) for ligand tethering studies^27^, whereas Nb_PTHR1-**X2**_ is uncharacterized in this context. Unlike experiments above, the binding sites of these Nbs are not defined by the location of engineered tags and they have not been determined experimentally. To elucidate the topology of conjugate-receptor interactions we sought to characterize the epitopes of Nb_PTHR1_ and Nb_PTHR1-**X2**_ on the receptor.

We labeled Nb_PTHR1_ and Nb_PTHR1-**X2**_ with detection tags via sortagging and analyzed binding to hPTHR1 stably expressed on HEK293 cells with flow cytometry. Both Nbs bound to hPTHR1 but only Nb_PTHR1_ bound to PTHR1-6E, suggesting the binding site of Nb_PTHR1-**X2**_ lies within the region of PTHR1 replaced by the 6E epitope, found within the disordered exon 2-encoded region of the receptor (Figure 4B, Supporting Figure 3A). This finding was corroborated with signaling assays in cells expressing PTHR1-6E (Supporting Figure 3) and rat PTHR1^36^ (Supporting Figure 4) on which Nb_PTHR1-**X2**_ conjugates were inactive. Addition of an exogenous synthetic peptide corresponding to the putative epitope of Nb_PTHR**-X2**_ blocked engagement of receptor by Nb_PTHR1-**X2**_ but not Nb_PTHR1_ (Supporting Figure 5). A binding assay performed with Nb_PTHR1_ and Nb_PTHR1-**X2**_ equipped with distinct labels demonstrated double labeling, further indicating that Nb_PTHR1_ and Nb_PTHR1-**X2**_ bind to separate epitopes (Supporting Figure 6).

**Figure 4:**
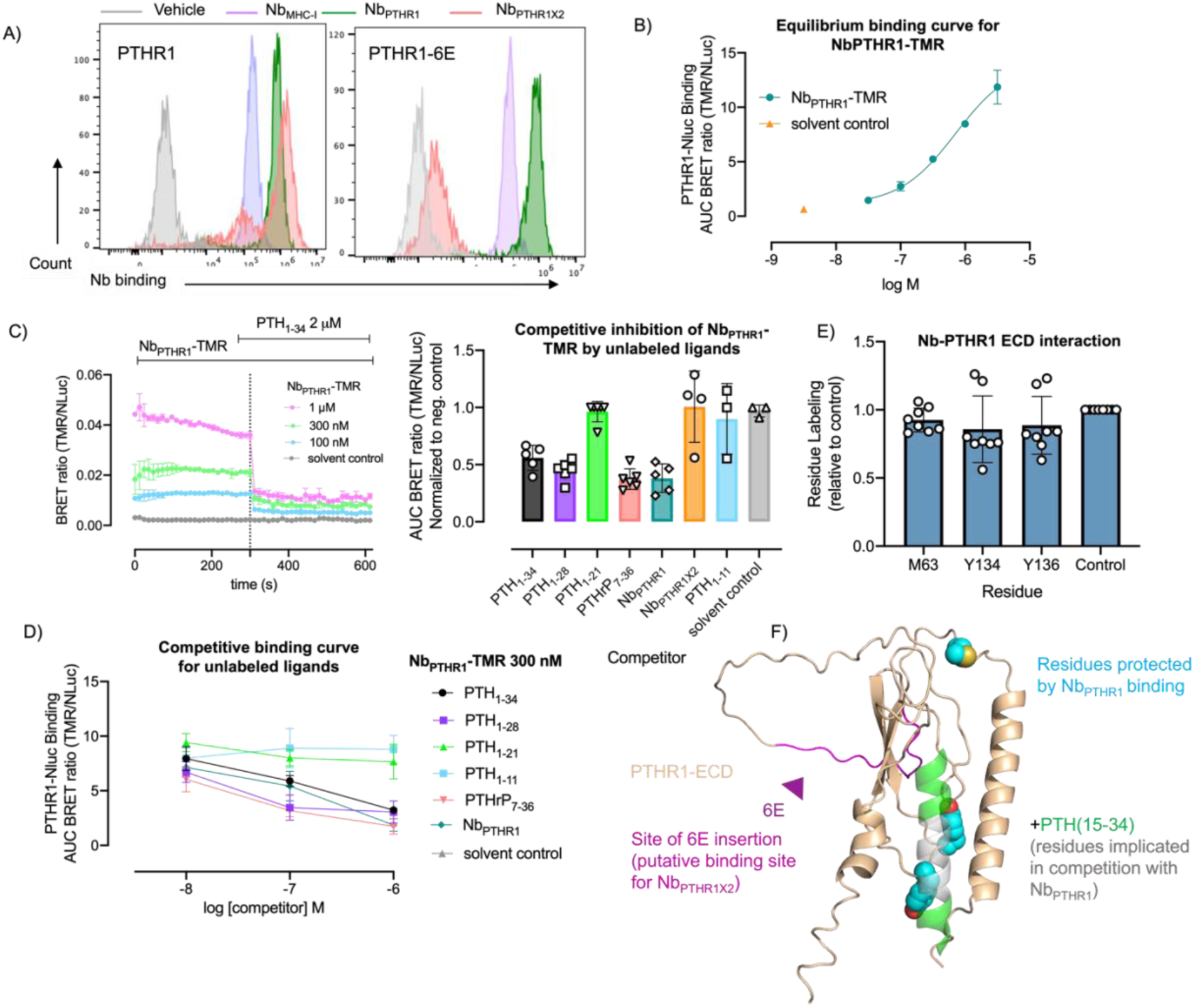
Evaluation of Nb_PTHR1_ and Nb_PTHR1X2_ binding to WT-PTHR1. A) Representative histograms for flow cytometry analysis of Nb_PTHR1_ and Nb_PTHR1-X2_ binding to PTHR1 and engineered receptors. Nbs (500 nM) labeled with biotin were incubated with cells expressing PTHR1 or PTHR1-6E, followed by washing, detection with streptavidin-APC, and assessment of cellular fluorescence. Analogous data for PTHR1-Nb_6E_ staining can be found in Supporting Figure 3. B) Representative concentration-response curve for BRET measurements of Nb_PTHR1_-TMR binding to cells expressing nLuc-PTHR1. Data points correspond to mean ± SD of AUC measurements, fit to a three-parameter logistic sigmoidal model. Measurements were performed in three independent experiments (Supporting Figure 7). C) Representative BRET measurements of Nb binding observed upon application of varying concentrations of Nb_PTHR1_-TMR followed by addition of unlabeled competitor peptide (PTH_1-34_, 2 µM). The bar graph shows summarized data for the inhibition of Nb_PTHR1_-TMR (1 µM) binding by unlabeled ligands. Data points correspond to mean ± SEM from 3-6 independent experiments, with quantitation as described in methods. See methods for peptide sequences. D) Concentration-response competition binding assays were performed using varying concentrations of unlabeled ligand added simultaneously with Nb_PTHR1_-TMR (300 nM). Data points correspond to mean ± SEM from 3 independent experiments. E) Analysis of Nb_PTHR1_-PTHR1 ECD interactions using hydroxy radical-based foot printing analysis (see Methods). Only the residues that show a trend towards protection from radical labeling upon addition of Nb_PTHR1_ are shown here with full data shown in Supporting Figure 11. Data points correspond to 8 independent replicates ± SD. Labeling is normalized to a control performed in the presence of a non-binding Nb. F) Alphafold2 model of PTHR1 ECD bound to PTH_15-34_ showing summarized Nb binding site characterization. The structure of full length ECD containing exon 2 (wheat) was generated from Alphafold2. This structure was aligned with the PTH_15-34_+ECD structure (PDB: 3C4M) using the “Align” command in Pymol.

To further define the binding site of Nb_PTHR1_, we used a BRET-based binding assay^37^. The affinity of tetramethylrhodamine (TMR)-labeled Nb_PTHR1_ for nanoluciferase-PTHR1 fusion (nLuc-PTHR1), in which nLuc was fused to the receptor N-terminus, was measured using BRET. Nb_PTHR1_-TMR exhibited modest affinity but robust labeling (Figure 4C, Supporting Figure 7), while PTH_1-34_-TMR showed stronger binding (Supporting Figure 8). We developed a competition binding assay on cells expressing nLuc-PTHR1 with Nb_PTHR1_-TMR and unlabeled competitor ligands (Figures 4C-D, Supporting Figure 9). We found that PTH_1-34_, PTH_1-28_, and PTHrP_7-36_ competed with TMR-Nb_PTHR1_ for binding, whereas more truncated ligands such as PTH_1-21_ did not, suggesting that the binding site of Nb_PTHR1_ overlaps with that of residues 21-28 of PTH_1-34_. Nb_PTHR1-**X2**_ also failed to compete Nb_PTHR1_-TMR for binding, providing further support of their separate binding sites (Supporting Figures 9 and 10).

These findings were corroborated by analysis of the interaction of Nb_PTHR1_ with purified PTHR1 extracellular domain (ECD) using hydroxy radical-based foot printing analysis based on plasma induced modification of biomolecules (PLIMB)^38^. PTHR1 ECD was expressed, refolded and purified, according to past work (see Supporting Figure 21 for the protein sequence) ^20^. Incubation of PTHR1 ECD with Nb_PTHR1_ suggested blockade of modification of ECD at three residues (Figure 4E, Supporting Figure 11), two of which (Y134, Y136) are found near the binding site of residues 20-24 of PTH in the PTH-PTHR1 ECD structure (Figure 4F)^39^. Although these differences in hydroxy radical labeling did not reach statistical significance (*p*∼0.10), they conform with predictions made from competition binding assays. Together, these data support a model in which Nb_PTHR1_ and Nb_PTHR1-**X2**_ bind at distinct sites on PTHR1 ECD, with the binding site of Nb_PTHR1_ overlapping in part with the binding site of residues 20-28 of PTH_1-34_.

Conjugates produced from Nb_PTHR1_ and Nb_PTHR1-**X2**_ with PTH_1-11_ were tested on cells expressing hPTHR1 in the panel of assays described for engineered receptors above (Figure 5A). In Gαs-cAMP assay, the conjugation of PTH_1-11_ to Nb_PTHR1_ or Nb_PTHR1-**X2**_ increased its potency by 10- to 100-fold (Fig. 5B). Additionally, both Nb_PTHR1_- and Nb_PTHR1-**X2**_-PTH_1-11_ conjugates induced substantially longer durations of cAMP production relative to PTH_1-11_ (Fig. 5C-D). Consistent with the ability of PTH_1-11_-Nb_PTHR_ conjugates to activate PTHR1 signaling through the Gαs pathway, they promoted Gαs dissociation from the plasma membrane assay with potencies that mirrored those of the cAMP assay (Table 1, Supporting Figure 12). Analogously to experiments with engineered receptors, both Nb_PTHR1_- and Nb_PTHR1-**X2**_-PTH_1-11_ conjugates displayed negligible recruitment of β-arrestin 2 to the plasma membrane or early endosomes in cells stably expressing hPTHR1 (Fig. 5D-E, Supporting Figure 12). To quantitatively compare the differences observed between the two signaling pathways, the bias model of agonism was used to calculate ΔΔLog (Emax/EC_50_) values (Supporting Figure 13). Only a crude estimate of bias factors was possible due to the weak activity of conjugates for inducing arrestin recruitment. We also found that Nb_PTHR1_-PTH_1-11_ failed to induce intracellular calcium mobilization (Fig. 5F), in line with findings from engineered receptors above.

**Figure 5:**
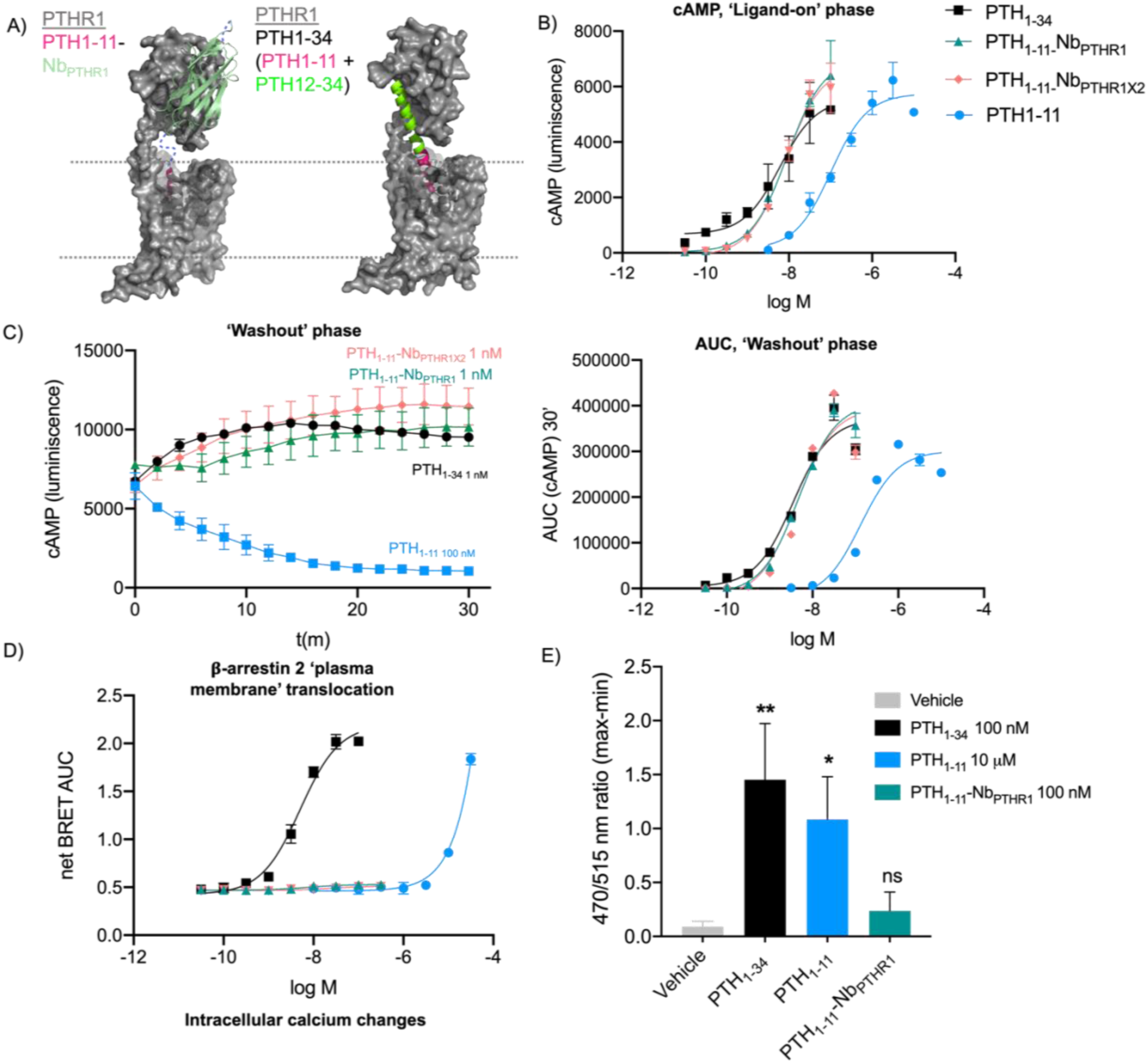
Evaluation of Nb-ligand conjugate signaling at WT-PTHR1. A) A schematic comparing a hypothetical mode of association between PTH_1-11_-Nb_PTHR1_ and PTHR1 with that of an experimental PTH_1-34_-PTHR1 complex (PDB: 6FJ3). Note that this model is for schematic purposes only and is not intended to model the site of Nb_PTHR1_ binding. B) Representative concentration-response curves for induction of cAMP responses in cells expressing PTHR1. C) (Left) Representative kinetic evaluations of ligand-induced responses following washout. (Right) Washout signaling was quantified as AUC and characterized in concentration-response plots. D) Representative concentration-response curves for ligand-induced recruitment of β-arrestin 2 to the plasma membrane. Endosomal β-arrestin 2 data is shown in Supporting Figure 12. For panels A-C data points are shown as mean ± SD from technical replicates in a single representative experiment, fit to a three-parameter logistic sigmoidal model. Each assay was evaluated in 3-5 biological replicates. Tabulation of agonist potency and summarized AUC washout dose-response parameters are shown in Table 1. E) Measurement of intracellular Ca^2+^ mobilization in response to indicated compounds. Data points correspond to means ± SEM from 3 independent experiments conducted with six technical replicates per experiment. Statistical significance was assessed by one-way ANOVA, with Dunnett’s post hoc correction (\**p* < 0.05; \*\**p* < 0.01; \*\*\**p* < 0.001; \*\*\*\**p* < 0.0001; ns not significant).

These observations prompted us to test whether Nb_PTHR1_-PTH_1-11_ induced receptor internalization. Observation of receptor trafficking by fluorescence microscopy showed that Nb_PTHR1_-PTH_1-11_ was inefficient in inducing receptor redistribution into intracellular puncta relative to PTH_1-34_ (Supporting Fig. 14). Results from microscopy experiments were corroborated by analysis with ELISA in which levels of receptor at the cell surface were measured following exposure to Nb_PTHR1_-PTH_1-11_ or PTH_1-34_ (Supporting Fig. 15). PTH_1-34_ caused a significant reduction in levels of cell surface PTHR1 whereas Nb_PTHR1_-PTH_1-11_ did not.

To test whether Nb_PTHR1_ binding alone caused variation in ligand signaling properties, we added Nb_PTHR1_ and peptide ligands separately (Supporting Figure 16), which demonstrated that the presence of Nb_PTHR1_ had a negligible impact on PTH_1-11_ or PTH_1-34_ signaling and bias. To assess the relative contributions of Nb and PTH_1-11_ for the affinity of these conjugates for PTHR1 we compared the binding of labeled Nb with labeled Nb-PTH_1-11_ conjugates (Supporting Figure 17). Nb_PTHR1_ and Nb_PTHR1-**X2**_ stained cells with similar intensity and potency when compared to their Nb-PTH_1-11_ conjugates, indicating that the Nb-receptor epitope interaction provides most of the binding affinity for these conjugates.

One hypothesis on why PTH_1-11_-Nb conjugates exhibit biased agonism is that they may engage with receptor in a manner that is qualitatively different from conventional ligands such as PTH_1-34_. Although PTH_1-34_ is comprised of two functional peptide domains that interact with different portions of the receptor, structural studies have shown this peptide binds to and acts upon a single receptor. We wondered whether PTH_1-11_-Nb conjugates could act by bridging two separate receptor protomers (“activation *in trans”*, Figure 6A). In this scenario, the Nb would bind to one receptor and the linked PTH_1-11_ would activate a different nearby receptor. Notably, there is evidence that receptor dimerization and oligomerization can impart biased signaling properties for class B GPCRs^40^. We sought to probe this hypothesis by extending the length of the linker between the binding and receptor activation elements of the PTH-Nb conjugates. A longer linker could help alleviate any signaling deficiency related to a “tug of war” scenario (Figure 6A, Right) or facilitate activation *in trans*. Even with extended linkers we observed little variation in signaling properties, including biased agonism (Supporting Figure 18). To provide a more direct assessment of activation *in trans* we transfected cells to express two distinct constructs of PTHR1: an N-terminally truncated derivative of PTHR1 (YFP-delNT-PTHR1) that is bound poorly by Nb_PTHR1_ but is signaling competent^27^ and full-length rat PTHR1 with mutations in its transmembrane portion (rPTHR1-null, R233Q/Q451K) that renders it signaling incompetent^36^ but is still recognized by Nb_PTHR1_ (Figure 6B). Nb_PTHR1_-PTH_1-11_ is weakly active on cells expressing YFP-delNT-PTHR1, but upon co-transfection with rPTHR1-null its biological activity improved substantially (Figure 6B-D). PTH_1-34_, which is also binds rPTHR1-null exhibited no such enhancement in activity upon co-transfection. A set of control experiments confirmed that both PTH_1-34_ and Nb_PTHR1_-PTH_1-11_ are highly active on cells transfected with WT-rPTHR1 (Figure 6D). This finding offers evidence that Nb-PTH_1-11_ conjugates can efficiently engage in receptor activation *in trans*, whereas PTH_1-34_ cannot. This behavior correlates with biased agonism observed in other assays and may be mechanistically related.

**Figure 6:**
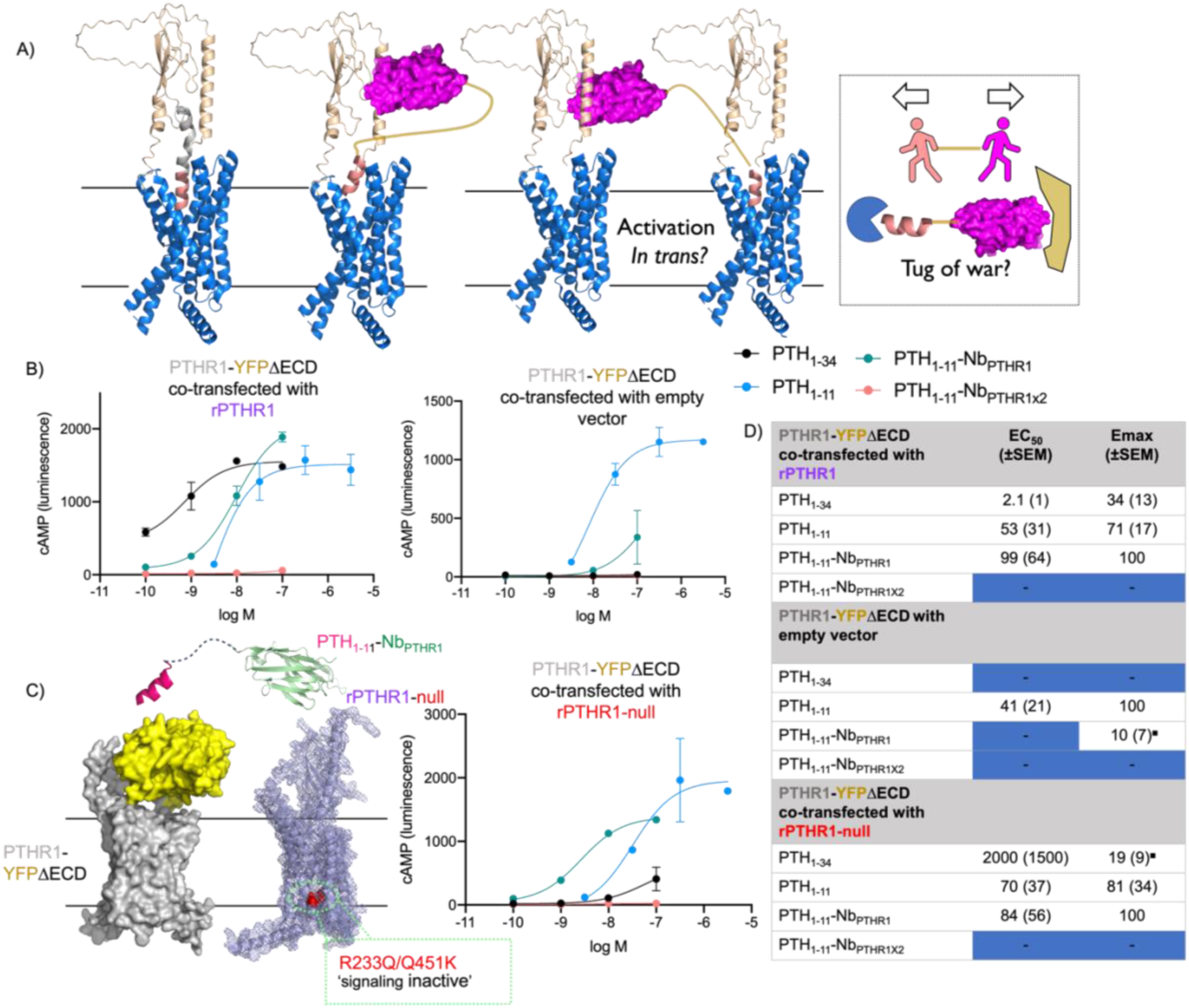
A) Proposed mechanisms to differentiate the binding and signaling of PTH_1-34_ (salmon and gray) and Nb-PTH_1-11_ (salmon and purple) to PTHR1 (blue and wheat). The standard model of receptor activation consists of a single ligand activating a single receptor (left). An alternative mechanism consists of one ligand acting upon two receptors in proximity “activation *in trans*” (middle). Signaling behavior might also relate to a discrepancy in linker length and the distance between binding sites for Nb_PTHR1_ and PTH_1-11_ in a “tug of war” type mechanism (right). B) Representative dose response data for induction of cAMP responses in cells co-transfected with PTHR1-YFPΔECD (PTHR1 ECD replaced with YFP) and other receptor plasmids. C) Schematic and data for PTH_1-11_-Nb_PTHR1_ activation *in trans* in a system co-expressing PTHR1-YFPΔECD and rPTHR1-null (R233Q/Q451K rPTHR1, signaling inactive). Schematic structures of PTHR1-YFPΔECD and rPTHR1-null were generated using Alphafold2 (see methods). Mutations in rPTHR1-null are highlighted in red. D) Tabulation of agonist potency and E_max_ parameters derived from 3 independent experiments shown as mean(±SEM). E_max_ values were normalized to the E_max_ of either PTH_1-11_-Nb_PTHR1_ or PTH_1-11_. ^▪^indicates that the value shown corresponds to the response recorded at the highest tested concentration rather than an E_max_ value generated by model fitting.

## Discussion

The prevailing view on how selected ligands manifest biased agonism is through stabilization of distinct receptor conformations (or conformational ensembles) that preferentially engage one intracellular coupling partner over another. Receptor conformational preferences are thought to be dictated by contacts engaged by the ligand binding to the orthosteric (or allosteric) site(s). In this study we showed that a ligand of PTHR1 (PTH_1-11_) that signals through all receptor engaged pathways, albeit with moderately diminished potency for β-arrestin recruitment, can be converted to a ligand that is highly biased for signaling through the Gαs pathway through linkage with receptor binding Nbs (or peptide). Such variations in the behavior between ligand and ligand-Nb conjugates are unexpected given that both PTH_1-11_ and PTH_1-11_-Nb conjugates contain the same core ligand structure, and differ only in whether they are linked to a receptor binding Nb. These observations are novel and expand the utility of the systematic approach for linking peptidic GPCR ligands with antibodies (or their fragments), known as CLAMP, that was only recently described^27^.

Our unexpected observation of PTH_1-11_-Nb bias poses the question of how and why pathway selective signaling occurs in this context. To rule out the possibility that binding of the tethering Nb or peptide was imparting an allosteric effect on receptor function we used a variety of approaches. We used engineered receptors in which we dictated the site of Nb or peptide tethering through introduction of an artificial binding site, either in the form of a genetically engrafted epitope tag or an engrafted nanobody. These engineered sites, placed in a non-conserved and disordered region of the receptor, are distant from the orthosteric site and any portion of PTHR1 thought to be involved in mediating its transition from an inactive to active state. Ligands conjugated to Nb or peptide that bound to these engineered sites were very highly biased in their signaling. For experiments with WT PTHR1, we characterized the receptor epitope to which two Nbs (Nb_PTHR1_ and Nb_PTHR1X2_) bound (Figure 4). The PTH_1-11_-Nb conjugate made from either Nb exhibited highly biased agonism at WT PTHR1, in line with results on engineered receptors. Addition of free Nb_PTHR1_ (not linked to PTH) also had no impact on ligand signaling and bias (Supporting Figure 16). These experiments collectively suggest that biased agonism results not from Nb binding *per se* but rather as an emergent property from linking PTH_1-11_ to an anchor that binds to the receptor at some ancillary location. This is contrasted with PTH_1-34_, in which PTH_1-11_ is anchored by the binding of the PTH_12-34_ fragment. Note, that PTH_1-34_ is more potent than PTH_1-11_ in all signaling pathways measured and does not exhibit strong ligand bias.

We hypothesize that variation in ligand anchoring mechanisms contribute to the high level of signaling bias observed for the PTH_1-11_-Nb conjugates. Two nonexclusive schemas can be used to conceptualize these effects. One possibility is that the topologic constraints imposed by Nb binding does not allow the simultaneous engagement of the receptor orthosteric site by the linked PTH_1-11_ (Figure 6A, “Tug of War”). This constraint might result in a transient or altered mode of binding of PTH_1-11_ at the receptor orthosteric site, potentially leading to ligand bias. This hypothesis is disfavored by the observation that variation of the linker length between Nb and PTH_1-11_ has little impact on the signaling properties observed (Supporting Figure 18). Another possibility is that Nb tethering facilitates receptor activation through a mode that involves two receptor protomers (activation *in trans*). We find preliminary evidence of a difference for receptor activation *in trans* in comparing PTH_1-34_ and PTH_1-11_-Nb_PTHR1_, which suggests this mechanism may be important for biased agonism, consistent with reports of GPCR assemblies facilitating biased agonism^40,41^. Whether Nb-ligand conjugation provides a generalizable approach for the design of biased agonists is under study. Notably, bitopic small molecule agonists have been shown to exhibit biased agonism at mu opioid receptor^42^; however, in this case both binding units associate with a single receptor protomer. Mounting evidence suggests that bitopic ligands that restrict GPCR conformational changes might be particularly rich sources of biased agonists^43^. Further insight into structural features of PTHR1 biased agonism is available from a recently reported structure of a Gαs-biased agonist (PCO371) bound to PTHR1^44^. It is possible that PTH1-11-Nb_PTHR1_ conjugates may elicit conformational changes similar to those seen in PCO371 bound receptor, although experimental confirmation is needed.

Efforts to apply Nb-ligand tethering to other receptors will require Nbs (or antibodies) that bind to the extracellular receptor face. At current, there are limited number of examples of Nbs that bind surface-exposed regions of GPCRs^45–50^, and only some of these Nbs have been structurally characterized^51–53^. New Nb screening technologies^54,55^ are emerging and will likely facilitate further progress. Another exciting possibility is that receptor activation *in trans* could operate across GPCR heteromers, or even assemblies incorporating plasma membrane-localized proteins that are not GPCRs. Caution is needed as past work has shown that activation *in trans* is not possible for all GPCR-membrane protein pairs expressed on the same cell surface^56^. Even with these caveats, the prospects of using straightforward conjugation methodology to generate highly active, highly biased ligands offers compelling prospects for future studies.

The development of compounds that target GPCRs and induce biased signaling responses is of substantial interest for therapeutic development. Biased ligands may serve to induce preferential activation of receptor-coupled pathways that promote therapeutic effects while minimizing signaling through pathways that mediate drug-related side effects^6^. Efforts to explore such applications are limited when suitably biased agonists are unavailable. Alternative methods, such as genetic knockout of GPCR signaling partners, offer insights into the consequences of biased agonism. For PTHR1, the deletion of β-arrestin 2 in mice prevented bone loss in response to continuous PTH stimulation, a side effect observed with conventional PTH-based therapeutics. This observation suggests that PTHR1 agonists biased towards Gαs might be candidates for osteoporosis therapeutics with reduced side effects. In this regard it is noteworthy, that the PTH_1-11_-Nb conjugates reported here appear to be the most highly Gαs-biased PTH agonist peptides reported to date. It is also worth noting that nanobody conjugation may be a general strategy to consider for delivering otherwise weak or unstable potential drug products to their intended target sites of action. Future efforts to generate biased agonists and related therapeutic lead candidates might benefit from the approach and principles outlined here.

## Methods

### Nanobody expression and purification

Nb protein sequences acquired from literature (previously named 22A3 and 23A3)^35^ were codon optimized for bacterial expression and cloned into a pET26b expression in frame with pelB and His6 sequences using clone EZ service from GenScript. The production and purification of Nb_6E_ (previously named VHH05) has been described previously^27^. Briefly, BL21(DE3) *E. coli* were transfected via heat shock with plasmids encoding nanobodies of interest and grown in medium (Terrific Broth) containing kanamycin (50 μg/mL). Transformed bacteria were used generate a starter culture, which was used to inoculate full-size cultures (1-4 L) containing kanamycin. This culture was grown at 37°C and expression induced with Isopropyl β-d-1-thiogalactopyranoside (IPTG, 1 mM) at an optical density (OD_600_) of 0.6. The induced culture was then shaken 30°C overnight.

Bacteria were harvested via centrifugation for 30 min at 6,000 RPM (Avanti J Series centrifuge) and resuspended in 30 mL of NTA wash buffer (tris buffered saline + 10 mM imidazole, pH 7.5) containing protease inhibitor (Pierce Protease Inhibitor Tablets, ThermoFisher A32953). Cells were then lysed via sonication and the lysate was centrifuged at 15,000 RPM for 45 min. After centrifugation, the Nb was purified from lysate by batch-based Ni-NTA chromatography, followed by size-exclusion chromatography (Cytiva Akta^TM^ / Pure) using a HiLoad^TM^ 16/600 Superdex 200 pg column with an isocratic gradient of TBS (Flow rate 1 mL/min). Fractions of interest were collected and concentrated using a 10kDa MW cutoff Amicon spin-concentrator. The identity of the purified fraction was confirmed by mass spectrometry, and the concentration of Nb was determined by measuring the absorbance at 280 nm. Aligned sequences of Nbs used in this study are shown in Supporting Figure 19.

### Peptide synthesis

All peptides were synthesized via solid phase peptide synthesis with Fmoc protection of the amine backbone on a Gyros PurePep Chorus or Liberty Blue Microwave-Assisted Automated Peptide Synthesizer. Where relevant, a Cys residue was incorporated at the C-terminus of the peptide for functionalization using Cys-maleimide chemistry (see below). Peptide synthesis was performed on Rink Amide resin (0.05 mmol scale) to afford a C-terminal carboxamide. Fmoc-amino acids were dissolved in dimethylformamide (DMF) and added to resin (8 equivalents) with HATU ((1 - [Bis(dimethylamino)methylene]-1H-1,2,3-triazolo[4,5-b]pyridinium 3-oxid hexafluorophosphate, 8 equivalents) and N,N-diisopropylethylamine (DIPEA, 16 equivalents). Fmoc groups were deprotected using 20% piperidine in DMF.

Upon completion of the synthesis, peptides were cleaved from the resin using a cleavage cocktail comprised of trifluoroacetic acid(TFA)/H_2_O/triisopropylsilane(TIS) (92.5:5:2.5% by volume) and rocked at room temperature for 3 hours prior to filtration. After the cleavage, the crude peptides were precipitated using chilled diethyl ether and pelleted by centrifugation (3,000 RPM for 2 minutes). Peptides were purified via preparative-scale HPLC using a Phenomenex Aeris Peptide XB-C18 Prep column (particle size 5 µM, 100 Å pore size) with a linear gradient of solvent A (0.1% TFA in H_2_O) and solvent B (0.1% TFA in acetonitrile). Fractions of interest were combined and lyophilized. Lyophilized peptides are then dissolved in DMSO at desired concentrations and frozen. Mass spectrometry characterization of peptides is shown in Supporting Table 1. Sequences for selected peptides used in competition binding assays is shown in Supporting Figure 20. The peptide sequences used in functional assays are different from those used in the competition binding assays.

PTH peptides with C terminal Cys residues purified by HPLC were incubated with 3 molar equivalents of DBCO-maleimide (Click Chemistry Tools #A108-100) and the DBCO-peptide products purified by HPLC. The purified product was lyophilized and dissolved in DMSO at a concentration of 1 mM. The identity of DBCO-modified PTH peptides was confirmed by LC-MS.

### Nb labeling via sortagging

Sortagging reactions were performed as previously described^27^ and were comprised of the following components: protein bearing a sortase recognition motif (LPETGG) followed by a hexa histidine tag at the C-terminus (20-200 µM final concentration), triglycine-probe conjugates (500-1000 µM final concentration), and Sortase 5M (10-20 µM final concentration). Reactions were performed in sortase buffer (10 mM CaCl_2_, 50 mM Tris, 150 mM NaCl, pH 7.5) and shaken at 12°C overnight. After incubation, the reaction was incubated with nickel NTA beads to capture Sortase 5M and unreacted starting protein. Uncaptured material was further purified using disposable desalting columns to remove triglycine conjugates (Cytiva PD-10 Sephadex^TM^ G-25M). Eluents were monitored for absorbance at the fluorescent probe absorbance wavelength (tetramethylrhodamine: 555 nm, Fluorescein: 494 nm), 220 nm, and 280 nm for the presence of protein conjugate. Fractions containing product were combined then concentrated by spin filtration (Amicon Ultra 0.5 mL Centrifugal Filters 10,000 NMWL).

### Click chemistry preparation of Nb-ligand conjugates

The peptide-DBCO conjugate was used in an azide-alkyne (“click”) reaction between the azide-functionalized Nb and a DBCO-modified synthetic PTH peptide (Figure 1), as previously described^27^. Nb-biotin-azide conjugates were mixed with an excess of PTH-DBCO (3-fold molar excess) in TBS. The reaction was shaken at 25 °C until unreacted Nb-biotin-azide has been completely consumed. The product conjugates were purified from free DBCO-modified peptide using a PD10 size exclusion column. Product identity was confirmed by LC-MS.

### Cell culture and receptor constructs

A Human Embryonic Kidney 293 cell line (HEK 293; ATCC CRL-1573) stably transfected with luciferase-based pGlosensor-22F cAMP reporter plasmid (Glosensor, Promega Corp) has been described previously [GS22^57^]. GS22 cells were used to generate cell line stably expressing either native human PTHR1, PTHR1-6E, or PTHR1-Nb_6E_ as previously described^27,29^. Plasmids encoding receptors under study have been previously defined ^27^. Sequences of all plasmids validated by Sanger sequencing. All cell lines were cultured in high-glucose DMEM (Gibco, Thermo Fisher Scientific), containing 10% fetal bovine serum (Sigma-Aldrich) and 1% of penicillin and streptomycin mixture and grown in an incubator at 37°C in a humidified atmosphere containing 5% CO_2_. The cells were passaged every 3–4 days and seeded to achieve a confluency of 60-70% for experiments relying on transient transfection. Cells plated for luciferase-based cAMP assays were grown in the same condition but were seeded to full confluency prior to assay execution. All cell lines were regularly checked for mycoplasma infection using the Lonza MycoAlert mycoplasma detection kit and were found to be negative. Aligned sequences for rat versus human PTHR1 are shown in Supporting Figure 21.

### Luminescence-based live cell cAMP accumulation assay

Cellular cAMP production was measured in living cells as described previously. In brief, cells in culture were trypsinized and transferred into clear bottom white-walled 96-well plates at a density of 80,000 cells per well. After achieving confluency, culture medium was removed and CO_2_ independent medium containing luciferin (0.5 mM) was added. Luminescence was measured until a stable background reading was obtained (∼ 10 min). Serial dilutions of ligands were then added such that the final well volume was 100 µL, with luminescence measured in real-time every 2 min for 12 min (Biotek Neo 2 plate reader). The peak luminescence responses (typically measured at 12 min) were used to generate concentration-response curves. The concentration response curves were fitted to individual experiments utilizing a sigmoidal dose-response model (log[agonist] vs. response [three parameters]; GraphPad Prism) to produce EC_50_ values. In instances where curves did not reach plateau at the highest concentration tested, curves were constrained to the Emax value observed for an index ligand run simultaneously.

The signaling duration of ligands was evaluated using “washout” assays in which the cells were stimulated with an agonist for a defined period as described above (ligand-on phase). After this period, the medium containing ligands was discarded. New CO_2_ independent medium containing fresh luciferin was added to all wells and luminescence responses was recorded for an additional 30 min (ligand-off phase). For competition-washout assays, the experimental protocol was as described above except that competitors were introduced during the ligand-off phase. Area under the curve (AUC) values, measured using GraphPad Prism, were used to construct dose-response curves for the washout assays.

For the dimerization luminescence-based cyclic AMP assays, HEK293 cells expressing a variant of PTHR1 with the extracellular domain replaced with yellow fluorescent protein (YFP) ^27^ were transfected with 1 µg of either HA-tagged wild-type rat PTHR1^36^, R233Q/Q451K rat PTHR1^36^, or no receptor. After transfection, the cells were grown and assayed for cyclic AMP production as described above.

### Flow cytometry analysis for Nb binding to receptors expressed in HEK293 cells

HEK293 cells stably expressing either native human PTHR1, PTHR1-6E, or PTHR1-Nb_6E_ were cultured as described above. Cells were harvested by trypsinization, which was quenched with the addition DMEM/FBS. Subsequently, the cells were transferred to a round bottom 96 well plate, pelleted by centrifugation (500 rpm for 3 min.), and resuspended in PBS containing 2% BSA (w/v) (PBS/BSA). Cells were incubated with varying concentrations of Nbs labeled with biotin using sortagging. Following incubation on ice for 30 min., cells were pelleted, washed, and resuspended in PBS/BSA containing streptavidin-APC (1:2000 dilution in PBS/BSA, (BioLegend #405207)) and incubated for 30 min on ice prior to washing. Washed cells were then resuspended in PBS/BSA for analysis by flow cytometry on a CytoFlex flow cytometer (Beckman Coulter). Intact cells were identified based on their forward scatter/side scatter profile and staining intensity was monitored in the APC channel (FL4). A minimum of 2,000 events corresponding to intact cells were recorded. Flow cytometry histograms were used to calculate median fluorescence intensity (MFI) values in each sample. MFI values were averaged among replicate samples.

### Bioluminescence resonance energy transfer (BRET) assays

#### Effector membrane translocation assay to measure β-arrestin and Gαs trafficking

HEK cell lines expressing receptors of interest were passaged and transfected as described above, with specific modifications based on a previously described protocol^34^. One day after seeding cells in a 10 cm dish, a transfection cocktail containing membrane-tethered BRET acceptor plasmid (rGFP-CAAX or rGFP-FYVE, 1008 ng), and RlucII-conjugated β-arrestin 2 BRET donor plasmid (β-arrestin 2-RlucII, 72 ng) and Lipofectamine 3000 was added following manufacturer instructions. For G-protein dissociation BRET assays, 1008 ng of rGFP-CAAX plasmid and 144 ng of Gαs-RlucII plasmid were used. Transfected cells were seeded at a density of 60,000 cells per well in a white 96-well plate and grown overnight. Prior to assays, culture medium was removed and replaced with Hanks Buffered Saline Solution (HBSS) supplemented with 5 mM HEPES. Cells were then treated with varying concentrations of ligand mixed with the luciferase substrate coelenterazine prolume purple (1 µM, NanoLight Technologies). The ligand concentration-dependent BRET change was measured using a Biotek Neo 2 plate reader. Signal at donor and acceptor wavelengths (410 nm and 515 nm, respectively) were measured every 150 seconds for a total of 30 min. BRET ratios (515/410 nm) were calculated and analyzed in GraphPad Prism v9. AUC values from kinetic measurements were used to generate concentration-response plots.

### Measurement of ligand binding kinetics using BRET

A cell line stably expressing a construct encoding nanoluciferase fused to the N-terminus of the PTHR1 was used as previously described^37^. This cell line was plated in black-walled 96-well plates as described above. On the day of the experiment, cell culture medium was removed and HBSS supplemented with 0.02% NaN_3_ and 5 mM HEPES was added (100 µl per well). The cells were then incubated in this buffer for at least 30 min at room temperature prior to the experiment. TMR-labeled peptides or Nbs were prepared at varying concentrations in HBSS/HEPES buffer containing 5 µM coelenterazine-h (NanoLight Technologies). Azide containing medium was removed from the assay plate and 100 µL of the peptide solution with coelenterazine-h was added to each well. Luminescence measurements were taken at 450 nm and 610 nm at 12 second intervals for 5 min (except for kinetic binding assays, described below).

In the washout (competition) assay mode, TMR-labeled Nb or peptide was first added at varying concentrations, followed by the addition of unlabeled ligands. Signals were recorded in two separate intervals: once when the labeled ligand was added alone, and once again after the unlabeled competitor was added. AUC of kinetic curves was determined to quantify changes in binding caused by competitor addition. Alternatively, in the pre-incubation competition assay format, varying concentrations of unlabeled ligands were added simultaneously with TMR-labeled Nb or peptide. In this assay format, the impact of competitor on labeled ligand binding was quantified by measuring BRET signal 12 minutes after addition.

For kinetic binding assays, BRET readings (at 450/610 nm) were taken in 7 second intervals for a total of 5 min following ligand addition. To measure the kinetics of ligand binding, the time course graphs were analyzed via the one-phase association method in GraphPad Prism, from which the k(obs) values were obtained (Equation 1) and plotted as a function of concentration onto a linear regression graph, which was used to derive the k(on) and k(off) values (Equation 2), which were, in turn, used to derive the k(D) value (Equation 3), as previously described^37^.

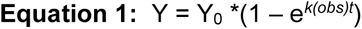

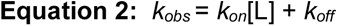

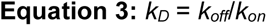

### Ligand induced intracellular calcium mobilization assays

The activity of various ligands for inducing intracellular calcium mobilization [Ca^2+^]_i_ was measured using Calbryte 520 AM (AAT Bioquest, USA) using the stably transfected cell lines described above. Briefly, 24h prior to the assay, cells were seeded into in 96-well black-walled plates (Thermo Fisher Scientific) and incubated overnight at 37°C. Calbryte 520 AM dye was dissolved in a HBSS in the presence of 0.04% Pluronic F-127, 20 mM HEPES, and 2.5 mM probenecid, which was added to each well and incubated at 37°C for at least 1 h before the assay. Dye loading solution was then removed and replaced with HBSS. Fluorescence was measured every 2 seconds (λ_excitation_ = 470 nm, λ_emission_ = 515 nm) using a high-throughput FLIPR^TETRA^ cellular screening system (Molecular Devices). Peptides were added after at least 2 min of background recording. Responses were quantified as relative fluorescence (“max-min”) calculated in ScreenWorks software (Molecular devices) and normalized to signal background.

### Hydroxy radical footprinting and analysis of nanobody-PTHR1 ECD interaction

Protein samples for analyzing Nb-PTHR1 ECD interactions were prepared, treated, and analyzed as previously described^38^ with some modifications. PTHR1 extracellular domain (ECD, see Supporting Figure 22 for sequence) was produced and purified as previously described ^20^. Nb_PTHR1_ or Nb_6E_ were incubated with PTHR1 ECD (1:1 molar ratio, 5.4 μM) at room temperature for 30 minutes prior to labeling. Samples were exposed to plasma induced modification of biomolecule (PLIMB) treatment conditions for 20 seconds. Following labeling, samples were quenched with a 5 *µ*L solution of 250 mM methionine in PBS (pH 7.4). Following PLIMB exposure, samples were proteolytically digested into peptides with trypsin. Samples were subjected to solid phase extraction using C18 StageTips and then analyzed using data-dependent acquisition with an Orbitrap Exploris 240 mass spectrometer.

The ‘.raw’ mass spectrometry data files were searched against the PTHR1 ECD sequence using the Protein Metrics Oxidative Footprinting Module. A list of standard expected modifications and expected PLIMB modifications was utilized in the database search. Peptides were identified using MS and MS/MS spectra, setting a 1% false discovery rate (FDR) cutoff. Changes in solvent accessibility for the trypsin digested samples were determined via comparison of the sum normalized intensities.

### Alphafold enabled receptor structural modeling

Structural models of the engineered receptors were generated through input protein sequences into an Alphafold2 Colab notebook^58^. The default AlphaFold2 settings were applied, which generates five models for each input sequence. Requisite models were applied to generate all graphics. Models were downloaded in PDB format and prepared as graphics using Pymol. In some cases, models were aligned with related experimentally determined structures using the “align” command in PyMOL.

### Data and statistical analysis

Data presented are either representative data from a single experiment (performed in technical duplicates, expressed as mean ± SD) or averaged (combined) data from at least three biological (independent) replicates (expressed as mean ± SEM). These distinctions are described in figure captions. Statistical analyses were performed only on collated data from biological replicates with n ≥ 3 using GraphPad Prism v9. Ligand bias (Fig. 4) was quantified by calculating the intrinsic relative activities of various ligands compared to a reference agonist (PTH_1-34_) using this equation^59^:

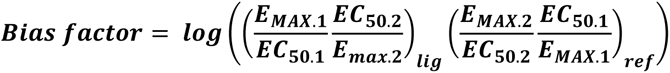

Parameters input into this equation were derived from curves fitted to graphs comprised of data from all biological replicates. In cases where ligands demonstrated no detectable response in specified assays, an estimated value for EC_50_ was generated from the dose-response curve described above.

Statistical significance (P<0.05) was assessed using Student’s *t* test (two-tailed), or one-way ANOVA as indicated in specific figure legends.

## Acknowledgements

We acknowledge the Massachusetts General Hospital peptide synthesis core facility (A. Khatri) for production of selected peptides. We thank M. Bouvier (University of Montreal) for provision of plasmids used for BRET-based assays of β-Arrestin and G-protein signaling. We thank Thomas Dean (Massachusetts General Hospital) for technical assistance with microscopy experiments. We acknowledge S. Gellman for provision of a construct encoding PTHR1 ECD and a cell line stably expressing nLuc PTHR1. We acknowledge the mass spectrometry core facility in NIDDK (J. Lloyd) for characterization of peptides and conjugates. This work was supported by the NIH Intramural Research Program (NIDDK, 1ZIADK075157, R.W.C.), PO1-DK11794 (T.J.G.), and funding from the NIH Director’s Award.

## Contributions

The paper was written by S.S. and R.W.C. Conceptual input was provided by S.S., T.G., and R.W.C. Experiments were designed and performed by S.S., B.C., T.G., and R.W.C.

---

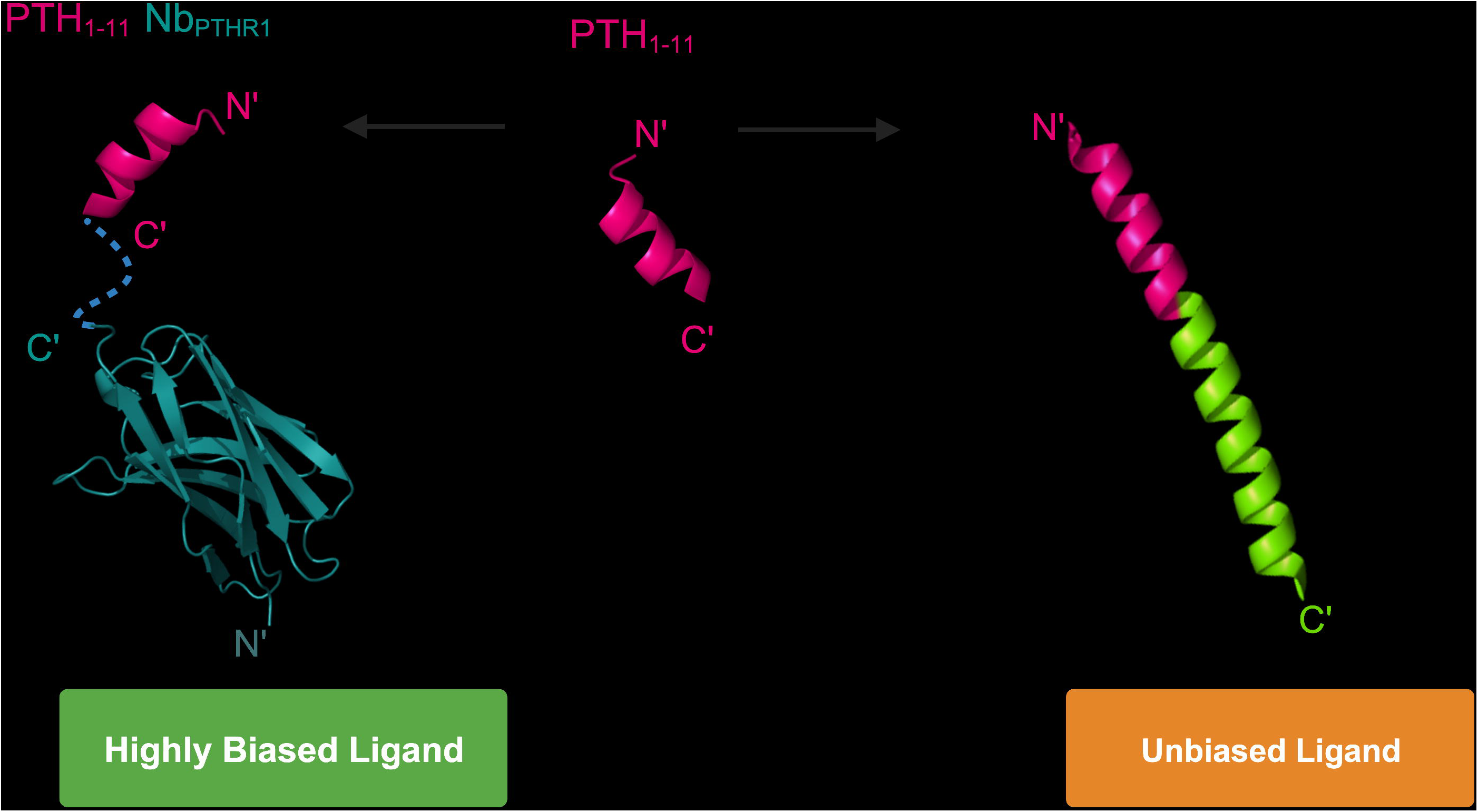

