## Supporting Information for "Highly biased agonism for GPCR ligands via nanobody tethering"

| Peptide | $[M+H]_{calc}$ | $[M+H]_{obs}$ |
| --- | --- | --- |
| $PTH_{1\text{-}34}$-Cys | 4218 | 4220 |
| $PTH_{1\text{-}11}$-Cys | 1412 | 1412 |
| $PTH_{1\text{-}11}$-Gly | 1366 | 1366 |
| $PTH_{1\text{-}11}$-DBCO | 1839 | 1839 |
| $PTH_{1\text{-}11}$-$PEG_4$-DBCO | 2086 | 2086 |
| $PTH_{1\text{-}21}$-Cys | 2667 | 2667 |
| $PTH_{1\text{-}11}$-6E | 3047 | 3046 |
| $G_3$-exon2 | 2175 | 2175 |
| $PTH_{1\text{-}11}$-$(Ahx)_2$-6E | 3160 | 3160 |
| $PTH_{1\text{-}11}$-Cys(TMR)-Ahx-azide | 2231 | 2231 |
| $PTH_{1\text{-}34}$-TMR | 4770 | 4771 |

**Supporting Table 1: Mass spectroscopy characterization of the peptides used in this study**. Peptides were analyzed by LC/MS as described in methods. Calculated masses ($[M+H]_{calc}$) refers to the monoisotopic mass of a singly protonated species. The masses recorded using mass spectrometry are labeled as $[M+H]_{obs}$.

| Conjugate | $[M+H]_{calc}$ | $[M+H]_{obs}$ |
|---|---|---|
| $Nb_{6E}$ | - | 13475 |
| $Nb_{6E}$-biotin-azide | 13221 | 13220 |
| $PTH_{1\text{-}11}$-$Nb_{6E}$ | 15164 | 15165 |
| $Nb_{PTHR1}$ | - | 14467 |
| $Nb_{PTHR1}$-DBCO | 14230 | 14228 |
| $Nb_{PTHR1}$-biotin-azide | 14207 | 14210 |
| $PTH_{1\text{-}11}$-$Nb_{PTHR1}$ | 16047 | 16051 |
| $Nb_{PTHR1}$-TMR | 14355 | 14356 |
| $PTH_{1\text{-}11}$-$Nb_{PTHR1}$-TMR | 16461 | 16464 |
| $Nb_{PTHR1X2}$ | - | 14703 |
| $Nb_{PTHR1X2}$-TMR | 14593 | |
| $Nb_{PTHR1X2}$-biotin-azide | 14443 | 14442 |
| $PTH_{1\text{-}11}$-$Nb_{PTHR1x2}$ | 16283 | 16284 |
| $Nb_{GFP}$ | - | 14232 |
| $Nb_{GFP}$-biotin-azide | 13978 | 13976 |
| $Nb_{MHC\text{-}I}$ | - | 14447 |
| $Nb_{MHC\text{-}I}$-biotin-azide | 14189 | 14188 |
| $Nb_{neg}$ (BC-2[1]) | - | 14992 |
| $Nb_{neg}$-biotin-azide | 14846 | 14845 |
| $PTH_{1\text{-}11}$-$Nb_{neg}$ | 16683 | 16684 |
| $PTH_{1\text{-}11}$-$PEG_4$-$Nb_{PTHR1}$ | 16293 | 16298 |

**Supporting Table 2: Confirmation of VHH-peptide conjugate identity using mass spectrometry.** Nb-peptide conjugates were analyzed by LC/MS as described in Methods. $MW_{calc}$ refers to the calculated average molecular weight and $MW_{obs}$ refers to the molecular weight recorded by mass spectrometry following analysis using MaxENT for deconvolution. $Nb_{Neg}$ corresponds to a nanobody that binds to the BC2 epitope, used here as a negative control[1]. Unlike the other Nbs used in this study, $Nb_{neg}$ was site-specifically modified with $G_3$-biotin-Ahx-azide instead of G3-botin-azide during the sortagging reaction. Dashes correspond to unmodified nanobody molecular weights, for which only experimentally observed masses were used for further calculations.

| Independent experiment (N) | cAMP | Washout AUC | Gαs | β-arrestin 2 (plasma membrane) | β-arrestin 2 (endosome) |
|---|---|---|---|---|---|
| **PTHR1-6E** | | | | | |
| $PTH_{1\text{-}34}$ | 6 | 6 | 4 | 4 | 3 |
| $PTH_{1\text{-}11}$ | 6 | 6 | 4 | 4 | 3 |
| $PTH_{1\text{-}11}$-$Nb_{6E}$ | 6 | 6 | 4 | 4 | 3 |
| $PTH_{1\text{-}11}$-$Nb_{neg}$ | 6 | 6 | 4 | 4 | 3 |
| **PTHR1-$Nb_{6E}$** | | | | | |
| $PTH_{1\text{-}34}$ | 4 | 4 | 3 | 6 | 3 |
| $PTH_{1\text{-}11}$ | 4 | 4 | 3 | 6 | 3 |
| $PTH_{1\text{-}11}$-Ahx-6E | 4 | 4 | 3 | 6 | 3 |
| **PTHR1** | | | | | |
| $PTH_{1\text{-}34}$ | 5 | 5 | 3 | 6 | 3 |
| $PTH_{1\text{-}11}$ | 5 | 5 | 3 | 6 | 3 |
| $PTH_{1\text{-}11}$-$Nb_{PTHR1}$ | 5 | 5 | 3 | 6 | 3 |
| $PTH_{1\text{-}11}$-$Nb_{PTHR1\mathbf{X2}}$ | 3 | 3 | 0 | 3 | 0 |

**Supporting Table 3**: Quantity of independent replicate experiments performed for both ligands and conjugates for data shown in Table 1 in the main text.

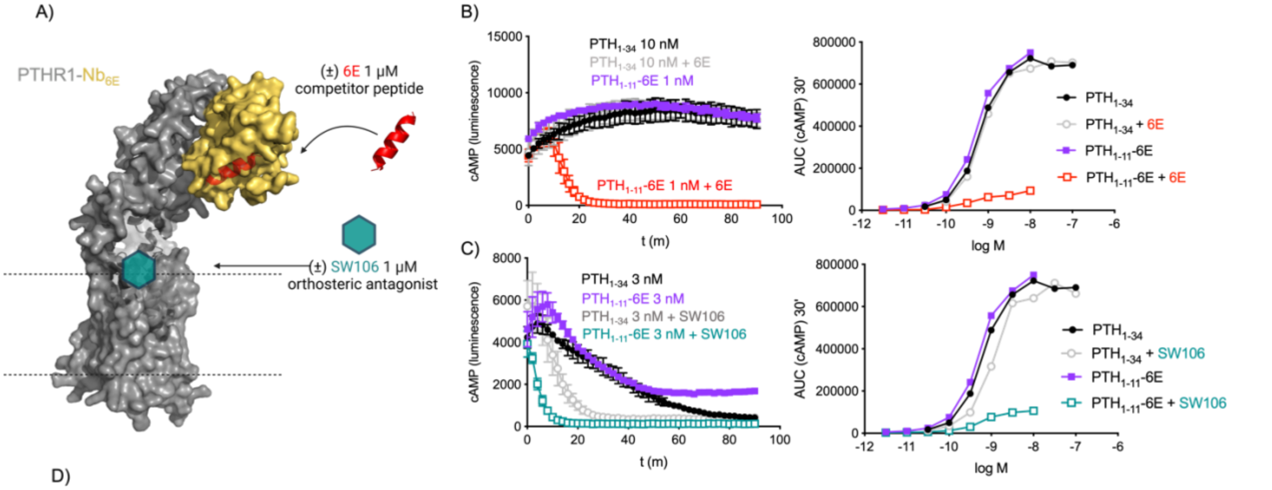


D)

| Ligands | cAMP, Washout AUC | | | | | |
|---|---|---|---|---|---|---|
| | Control | | +6E | | +SW106 | |
| | $EC_{50}$ (±SEM) | % $PTH_{1-34}$ activation $E_{max}$ (±SEM) | $EC_{50}$ (±SEM) | % $PTH_{1-34}$ activation $E_{max}$ (±SEM) | $EC_{50}$ (±SEM) | % $PTH_{1-34}$ activation $E_{max}$ (±SEM) |
| $PTH_{1-34}$ | 1.2 (0.7) | 100 | 0.7 (0.3) | 100 | 2.2 (3.7) | 100 |
| $PTH_{1-11}$-6E | 0.18 (0.12) | 110 (15) | - | 16 (6) | - | 13 (7) |

**Supporting Figure 1: Impact of antagonists on washout responses of ligand conjugates** A) Schematic of the washout competition assay for PTHR1-$Nb_{6E}$ in the continued presence of synthetic 6E peptide or orthosteric PTHR1 antagonist, SW106. B) Time course for cAMP production in HEK293 cells stably expressing PTHR1-$Nb_{6E}$ after washout of ligands in the presence or absence of competitor 6E peptide. Data sets in the right correspond to values generated from quantifying the area under the curve for kinetic washout responses. C) Analogous data for cAMP washout responses in the presence or absence of antagonist, SW106. Data points correspond to mean and associated SD from technical replicates. D) Compiled tabulation of agonist potency and Emax parameters derived from 3 independent experiments. For compounds where a plateau was not reached, Emax corresponds to the response observed at the highest dose. Data correspond to presented as mean (±SEM). A dash indicates activity was too weak to calculate an $EC_{50}$ value.

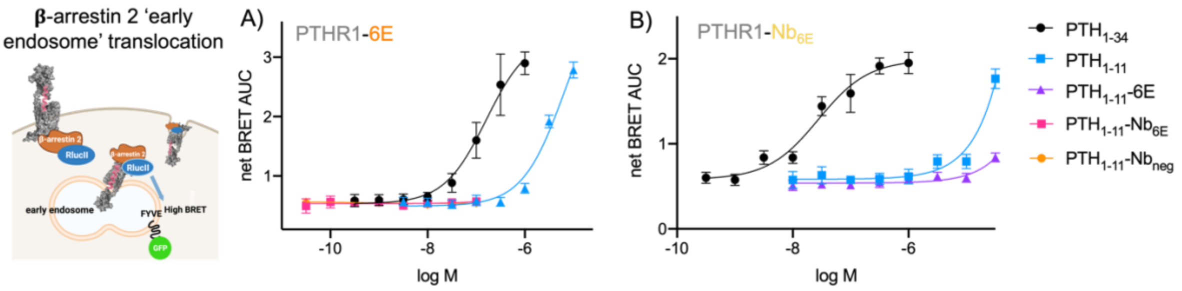


**Supporting Figure 2: Characterization of endosomal β-arrestin 2 recruitment by ligand conjugates**. A) Stimulation of β-arrestin 2 translocation to endosomes in HEK cells expressing PTHR1-6E. An increase in BRET ratio indicates ligand-induced β-arrestin 2 translocation to endosomes. Data are presented as AUC generated from BRET kinetic measurements. B) Analogous data for cells expressing PTHR1-$Nb_{6E}$. Data points correspond to mean and SD from technical replicates in a representative experiment, fit to a three-parameter logistic sigmoidal model. Tabulation of agonist potency parameters are shown in main Table 1, derived from 3-5 independent experiments.

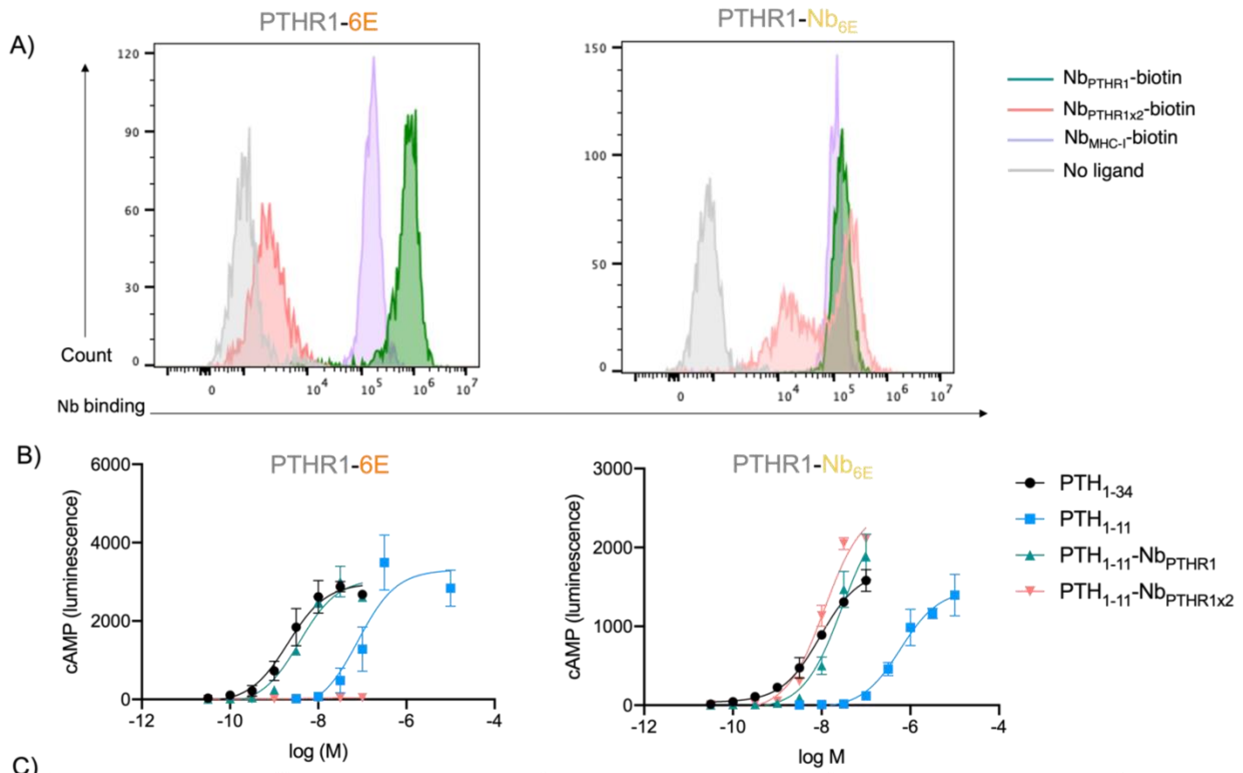


C)

| $EC_{50}$ (±SEM) Ligands | PTHR1 | PTHR1-6E | PTHR1-$Nb_{6E}$ |
|---|---|---|---|
| $PTH_{1\text{-}34}$ | 19 (16) | 4 (2) | 6 (2.5) |
| $PTH_{1\text{-}11}$ | 277 (214) | 94 (14) | 447 (173) |
| $PTH_{1\text{-}11}$-$Nb_{PTHR1}$ | 6 (3) | 3.4 (1.7) | 27 (13) |
| $PTH_{1\text{-}11}$-$Nb_{PTHR1X2}$ | 13 (7) | - | 6 (4.1) |

**Supporting Figure 3: Assessment of $Nb_{PTHR1}$ and $Nb_{PTHR1X2}$ acting on engineered receptors.** Representative histograms for flow cytometry analysis of $Nb_{PTHR1}$ and $Nb_{PTHR1\text{-}X2}$ binding to PTHR1-6E and PTHR1-$Nb_{6E}$ receptors. Nbs (500 nM) labeled with biotin were incubated with cells expressing PTHR1-6E or PTHR1-$Nb_{6E}$, followed by washing, detection with streptavidin-APC, and assessment of cellular fluorescence. No ligand refers to cells not exposed to biotin-labeled Nbs. B) Dose-response curves for maximal cAMP responses generated upon addition of ligands to cells expressing indicated receptors. Curves were generated by fitting to a 3-parameter logistic equation. C) Compiled tabulation of agonist potency parameters for ligand-conjugates. $EC_{50}$ values correspond to mean (±SEM) measurements from 3 or more assays run in duplicates. A dash indicates activity was too weak to calculate an $EC_{50}$ value. Note that a subset of this data (i.e. $EC_{50}$ for $PTH_{1\text{-}34}$ and $PTH_{1\text{-}11}$) are duplicated in the main text Table 1.

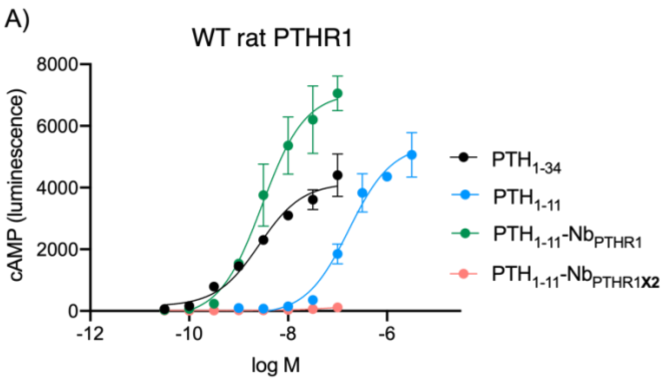


B)

| Ligands | Rat PTHR1, cAMP $EC_{50}$ (±SEM) |
|---|---|
| $PTH_{1\text{-}34}$ | 6.6 (3) |
| $PTH_{1\text{-}11}$ | 117 (40) |
| $PTH_{1\text{-}11}$-$Nb_{PTHR1}$ | 1.9 (0.7) |
| $PTH_{1\text{-}11}$-$Nb_{PTHR1X2}$ | - |

**Supporting Figure 4: Dose response data for Nb-ligand conjugates for cAMP production was assessed on cells expressing WT rat PTHR1.** Curves were generated using a three-parameter logistic sigmoidal model from a single representative experiment. B) Compiled tabulation of agonist potency parameters for ligand-conjugates at WT rat PTHR1. $EC_{50}$ values correspond to mean (±SEM) measurements from 3 independent biological replicates. A dash indicates activity was too weak to calculate an $EC_{50}$ value.

A)

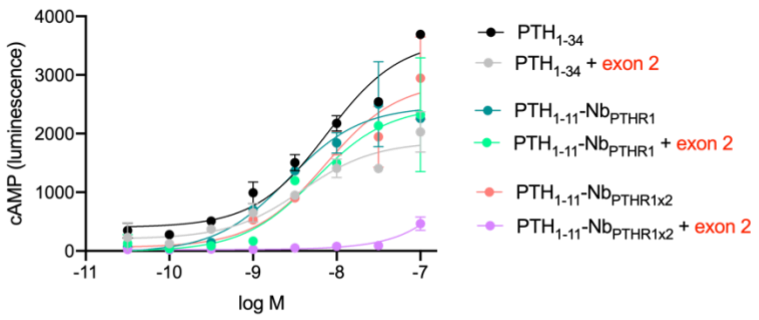


B)

| PTHR1 | $EC_{50}$, nM | SEM |
|---|---|---|
| without exon 2 | | |
| $PTH_{1\text{-}34}$ | 8 | 0.5 |
| $PTH_{1\text{-}11}$-$Nb_{PTHR1}$ | 4.3 | 2.2 |
| $PTH_{1\text{-}11}$-$Nb_{PTHR1X2}$ | 6.1 | 1.2 |
| with exon 2 | | |
| $PTH_{1\text{-}34}$ | 5.1 | 2.3 |
| $PTH_{1\text{-}11}$-$Nb_{PTHR1}$ | 4 | 1 |
| $PTH_{1\text{-}11}$-$Nb_{PTHR1X2}$ | >100 | - |

**Supporting Figure 5: Blockade of $Nb_{PTHR1\text{-}X2}$ binding through exogenous peptide derived from PTHR1 exon 2**. Dose response data for Nb-ligand conjugates for cAMP production was assessed on WT PTHR1 treated with synthetic exon 2 peptide (GGGWTSASTSGKPRKDKASGKL, 1.7 μM). This peptide encompasses the sequence of the portion of PTHR1 replaced by epitope tag in PTHR1-6E. Curves were generated using a three-parameter logistic sigmoidal model from a single representative experiment with or without exon 2 peptide. B) Compiled tabulation of agonist potency parameters for ligand-conjugates at PTHR1. $EC_{50}$ values correspond to mean (±SEM) measurements from 3 biological replicates. A dash indicates activity was too weak to calculate an $EC_{50}$ value.

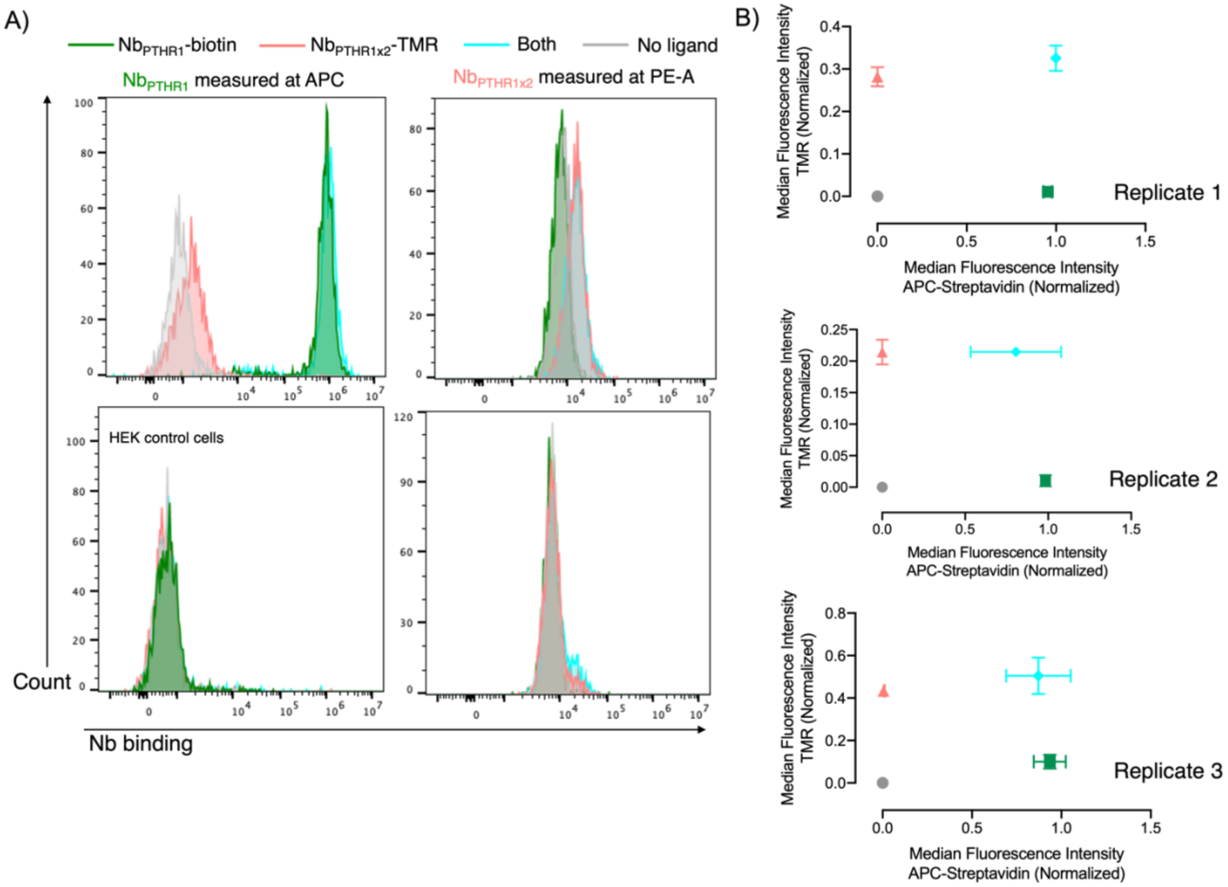


**Supporting Figure 6: Assessment of $Nb_{PTHR1}$ and $Nb_{PTHR1-X2}$ binding.** Representative histograms for flow cytometry analysis of $Nb_{PTHR1}$ and $Nb_{PTHR1-X2}$ with distinct labels binding to PTHR1. $Nb_{PTHR1}$ labeled with biotin (300 nM) and $Nb_{PTHR1-X2}$ labeled with TMR (300 nM) were incubated with cells expressing PTHR1, followed by washing, detection with streptavidin-APC, and measurement. “No ligand” refers to cells not exposed to labeled Nbs. “Both” refers to simultaneous treatment of cells with two distinctly labeled Nbs added together. The histogram in the bottom row of panels represents the same labeling experiment run on untransfected cells that do not express PTHR1. B) Quantified data normalized to the maximum signal observed for an index ligand presented as median fluorescence intensity values. Each plot corresponds to an independent experiment. Data points correspond to mean ± SD for technical replicates in each experiment.

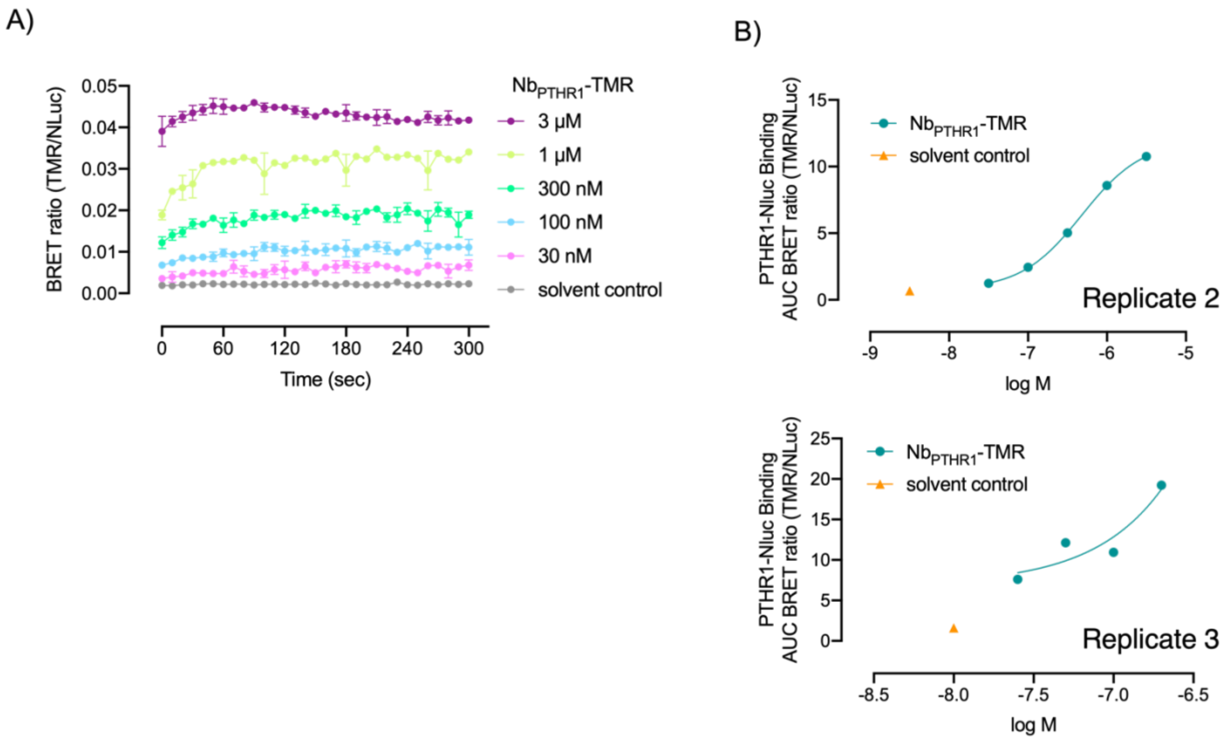


**Supporting Figure 7: Evaluating Nb$_{PTHR1}$ binding to PTHR1 using BRET.** A) Representative BRET kinetic measurement curve for the binding (association) of increasing concentrations of Nb$_{PTHR1}$-TMR to HEK cells expressing nLuc-PTHR1. Data are shown as mean ± SD from technical replicates in a single representative experiment. Note that by the time measurements begin the BRET signal is already at a plateau level, precluding calculation of association and dissociation kinetics. B) Concentration-response curves for biological replicates of BRET measurements of Nb$_{PTHR1}$-TMR binding to nLuc-PTHR1. Data points correspond to quantification of the area under the curve for kinetic association curves. Data are fitted to a 3-parameter sigmoidal dose response model. The third technical replicate is found in the main text (Figure 4B).

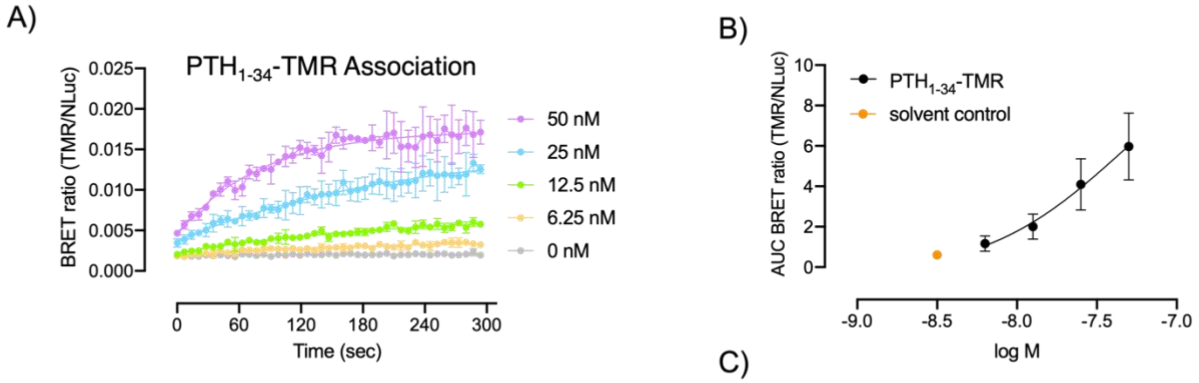


C)

| $PTH_{1-34}$ | |
|---|---|
| $K_{on}$ (x$10^5$ $s^{-1}M^{-1}$) | 3.6 ± 0.3 |
| $K_{off}$ ($s^{-1}$) | 0.003 ± 0.0002 |
| $K_d$ (nM) | 14 ± 1 |

**Supporting Figure 8: Evaluation of $PTH_{1-34}$ binding to PTHR1 using BRET.** A) Representative BRET kinetic measurements of the binding (association) of increasing concentrations of $PTH_{1-34}$-TMR (0-50 nM) added at time zero to HEK cells expressing nLuc-PTHR1. Data are shown as mean ± SD from technical replicates in a single representative experiment. B) Representative concentration-response curve for BRET measurements of $PTH_{1-34}$-TMR binding to nLuc-PTHR1. Data set generated from quantifying the area under the curve for kinetic association curve. Data are fitted to a 3-parameter sigmoidal dose response model. C) Tabulation of kinetic parameters and dissociation constants of $PTH_{1-34}$ binding to nLuc-PTHR1 fitted to a one-phase association model (see Methods). Kinetic parameters correspond to means ± SEM from 3 independent experiments.

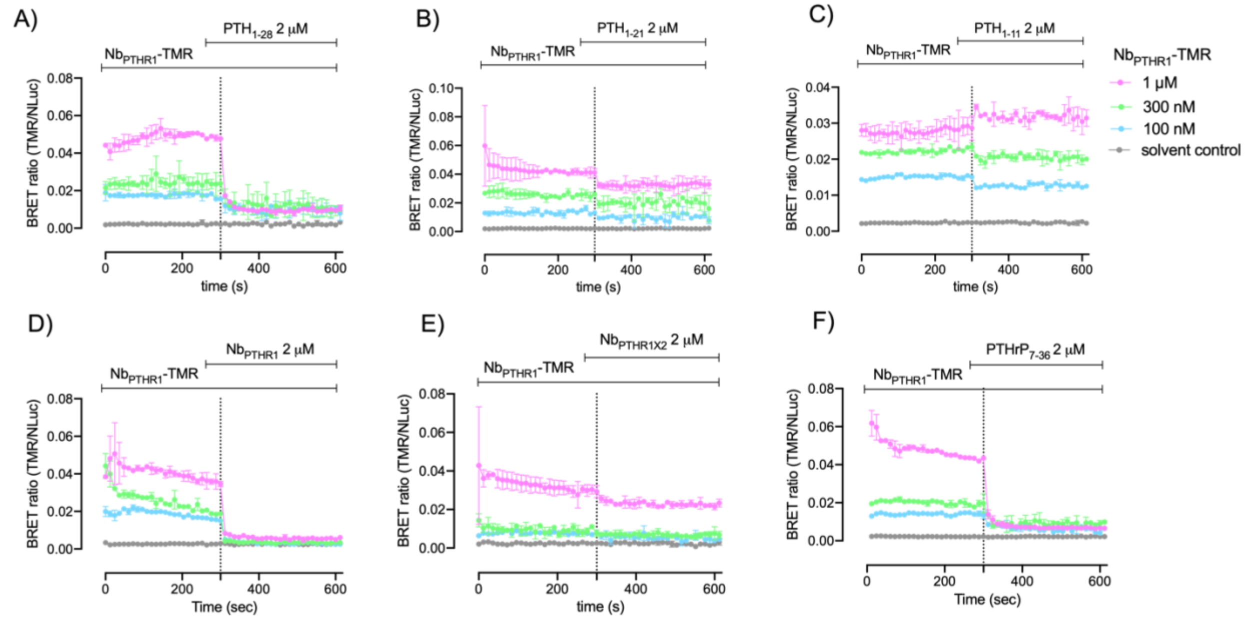


**Supporting Figure 9: Kinetic traces of BRET binding experiments for the competition of $Nb_{PTHR1}$-TMR with unlabeled competitors.** Representative BRET kinetic traces of Nb binding observed upon application of varying concentrations of $Nb_{PTHR1}$-TMR followed by addition of unlabeled ligand (A) $PTH_{1-28}$, (B) $PTH_{1-21}$, (C) $PTH_{1-11}$, (D) $Nb_{PTHR1}$, (E) $Nb_{PTHR1\text{-}X2}$, and (F) $PTHrP_{7-36}$ at a concentration of 2 µM. Quantified summaries for the these experiments is shown in Figure 4C. Data points correspond to mean ± SD from technical replicates in a single representative experiment.

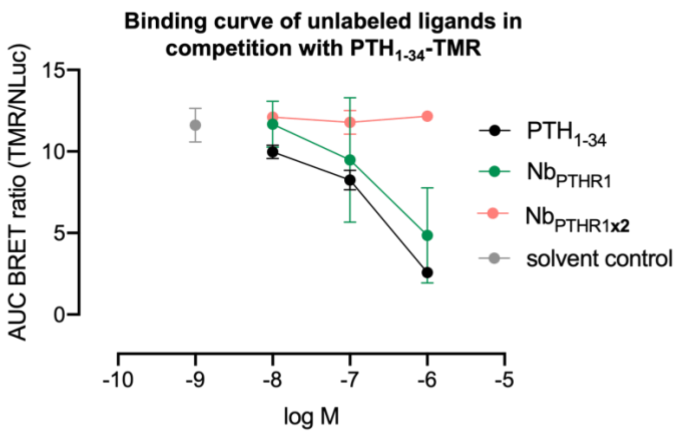


**Supporting Figure 10: Competition BRET binding assays with $PTH_{1-34}$-TMR.** Concentration-response competition binding assays were performed using varying concentration of unlabeled Nbs (or peptide) added simultaneously with $PTH_{1-34}$-TMR (300 nM). Data points correspond to mean (BRET AUC) ± SEM from 3 independent experiments.

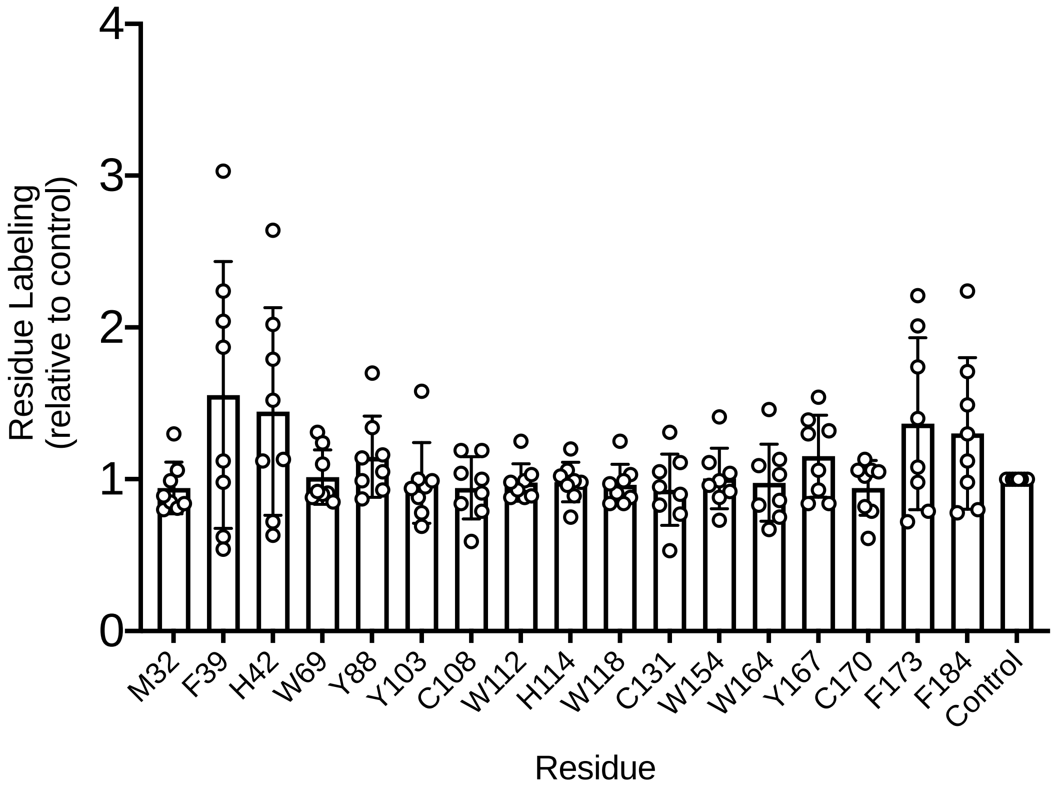


**Supporting Figure 11: Analysis of $Nb_{PTHR1}$-PTHR1 ECD interactions using hydroxy radical-based footprinting analysis**. Data points correspond to 8 independent replicates ± SD. Labeling is normalized to a control performed in the presence of a non-binding Nb ($Nb_{6E}$). See Figure 4E for residues showing a reduction in binding upon addition of $Nb_{PTHR1}$.

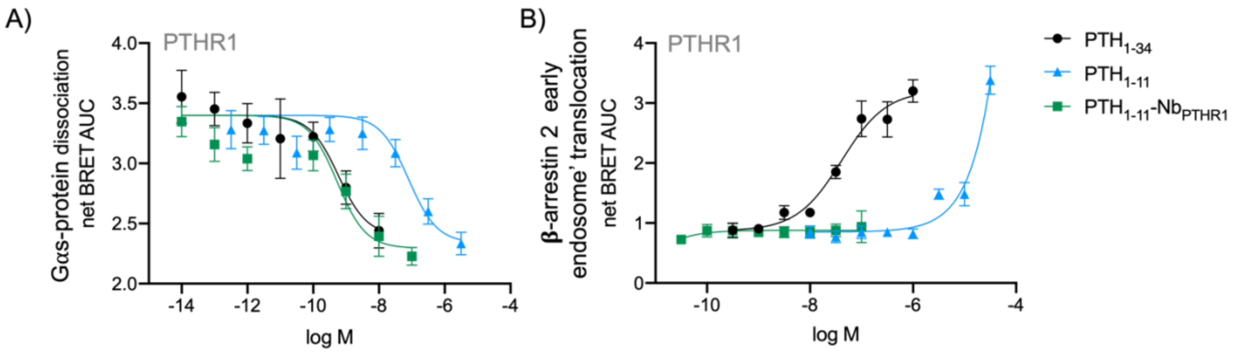


**Supporting Figure 12: Characterization of Nb-ligand conjugate signaling at WT PTHR1 through diverse pathways.** Concentration-response curves for ligands were measured for the A) dissociation of Gαs protein, and B) recruitment of β-arrestin 2 to endosomes in HEK cells expressing WT PTHR1. All responses are presented as AUC generated from kinetic BRET measurements. Data points correspond to mean ± SD from technical replicates in a representative experiment. Curves are fit to a three-parameter logistic sigmoidal model. Tabulation of agonist potency parameters are show in main Table 1, derived from 3-5 independent experiments (see Supporting Table 3).

| Ligand | G protein | | | | Arrestin | | | | Ligand bias | |
|---|---|---|---|---|---|---|---|---|---|---|
| | Emax (%) | $EC_{50}$ (nM) | Log ($Emax/EC_{50}$) | ΔLog ($Emax/EC_{50}$) | Emax (%) | $EC_{50}$ (nM) | Log ($Emax/EC_{50}$) | ΔLog ($Emax/EC_{50}$) | ΔΔLog ($Emax/EC_{50}$) | Bias factor |
| PTH1-34 | 100 | 6.1 | 8.2 | 0.0 (ref.) | 100 | 5.2 | 8.3 | 0.0 (ref.) | 0.0 | 1.0 |
| $Nb_{PTHR1}$-PTH1-11 | 100 | 9.2 | 8.0 | -0.2 | 100 | 1000000 | 3.0 | -5.3 | 5.1 | 127000 |
| PTH1-11 | 100 | 103 | 7.0 | -1.2 | 92 | 17230 | 4.7 | -3.6 | 2.3 | 213.3 |
| $Nb_{PTHR1X2}$-PTH-11 | 100 | 8.2 | 8.1 | -0.1 | 100 | 1000000 | 3.0 | -5.3 | 5.2 | 143000 |

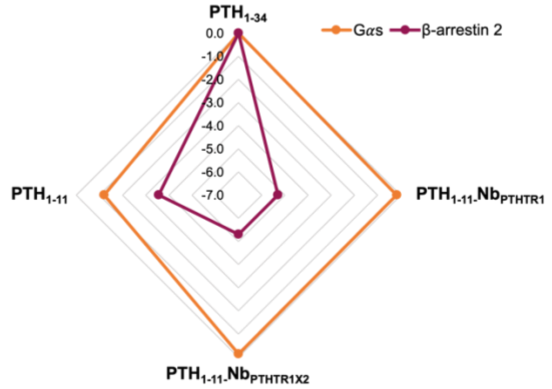


**Supporting Figure 13: Quantified bias factors for ligand-conjugates across functional assays.** Potency ($EC_{50}$) and efficacy (Emax, percent of $PTH_{1-34}$) values for Gαs-cAMP and β-arrestin 2 recruitment for PTHR1 were generated from compiled data fit to three parameter logistic equation. Since the Nb-conjugates were virtually inactive in the arrestin recruitment assay, top and bottom parameters were constrained to match those of $PTH_{1-34}$ to provide $EC_{50}$ values for ligand bias calculations. ΔLog($Emax/EC_{50}$) values were calculated relative to $PTH_{1-34}$ within each individual experiment. ΔΔLog($Emax/EC_{50}$) were then calculated between the indicated assays as described in Methods[2]. Bias factors were plotted as radar plot. The radar plot is based on the agonist type plotted against G protein and β-arrestin signaling. The values plotted correspond to ΔLog($Emax/EC_{50}$) values for each pathway.

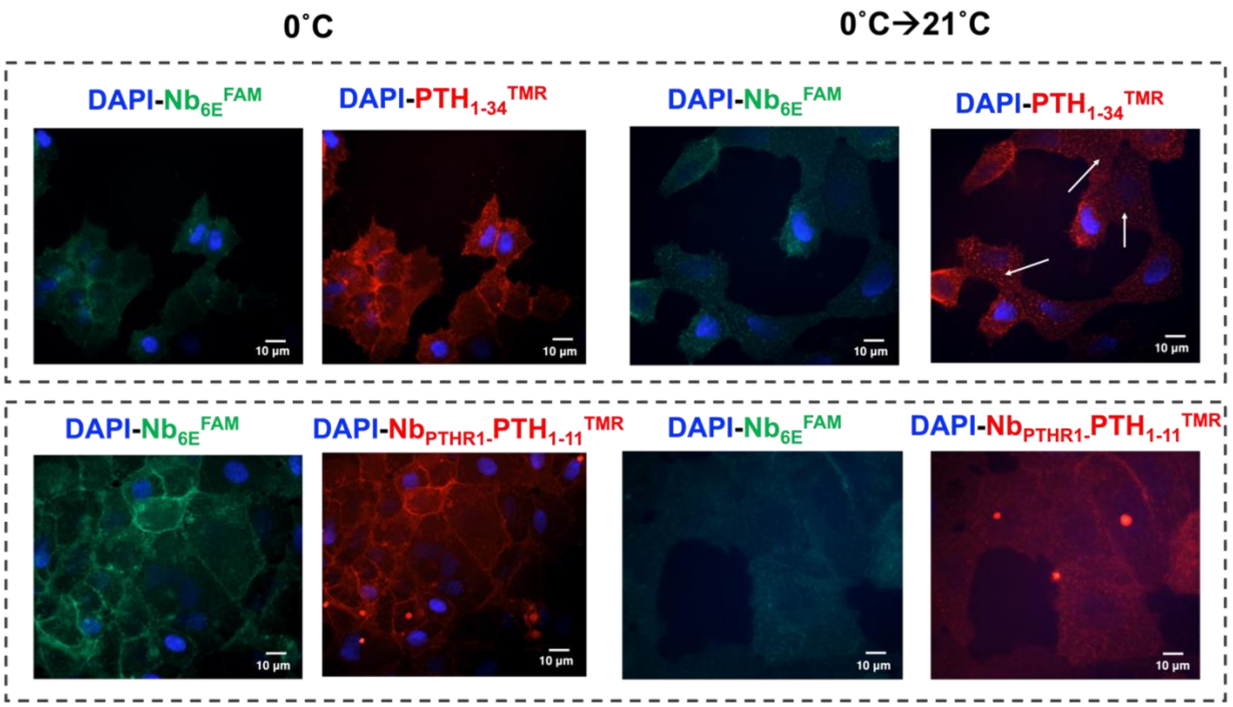


**Supporting Figure 14: Fluorescence microscopy analysis of ligand-induced internalization of PTHR1-6E.** HEK cells expressing PTHR1-6E were visualized through staining with Nb$_{6E}^{FAM}$ (green), DAPI (blue), and TMR-labeled ligand (red). Shown are representative images acquired after 30 min of stimulation with either 300 nM of PTH$_{1-34}^{TMR}$ (top) or PTH$_{1-11}$-Nb$_{PTHR1}^{TMR}$ (bottom). PTH$_{1-11}$-Nb$_{PTHR1}^{TMR}$ was synthesized from PTH$_{1-11}$-Cys(TMR)-Ahx-Azide and Nb$_{PTHR1}$-DBCO. Staining was performed on ice and incubation was carried out at either at 0°C (left) or 21°C (right). Following incubation cells were exposed to fixative prior to imaging. Scale bars, 10 $\mu$m. White arrows indicate punctate signals corresponding to ligand-induced intracellular accumulation of receptors.

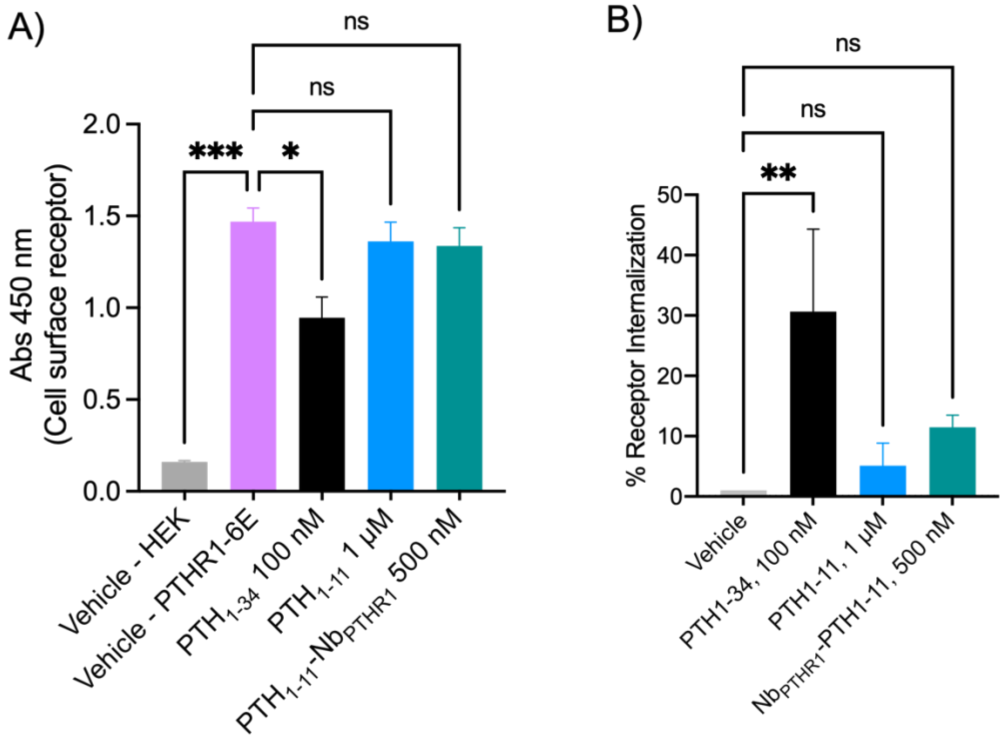


**Supporting Figure 15: Whole-cell ELISA to measure changes in surface levels of PTHR1-6E upon ligand exposure.** A) Intact, adherent cells expressing PTHR1-6E were exposed to ligand or vehicle at 37°C. After incubation, cell surface receptors were detected with $Nb_{6E}$-biotin and secondary staining with horseradish peroxidase-conjugated streptavidin. A) Representative data showing signal from detection of cell surface receptor following 30 minutes of treatment with indicated ligands. "Vehicle-HEK" indicates the use of cells not expressing PTHR1-6E and without exposure to ligands. "Vehicle-PTHR1-6E" indicates the use of cells expressing PTHR1-6E without exposure ligands. B) Graphic showing receptor internalization, quantified as the percent loss of cell surface receptors in ligand-treated cells normalized to Vehicle-HEK (negative control cells). Data points correspond to mean ± SEM from 3 independent experiments conducted with technical replicates in each independent experiment. Statistical significance was assessed by one-way ANOVA, with Dunnett's post hoc correction (*$p < 0.05$; **$p < 0.01$; ***$p < 0.001$; ****$p < 0.0001$; ns not significant).

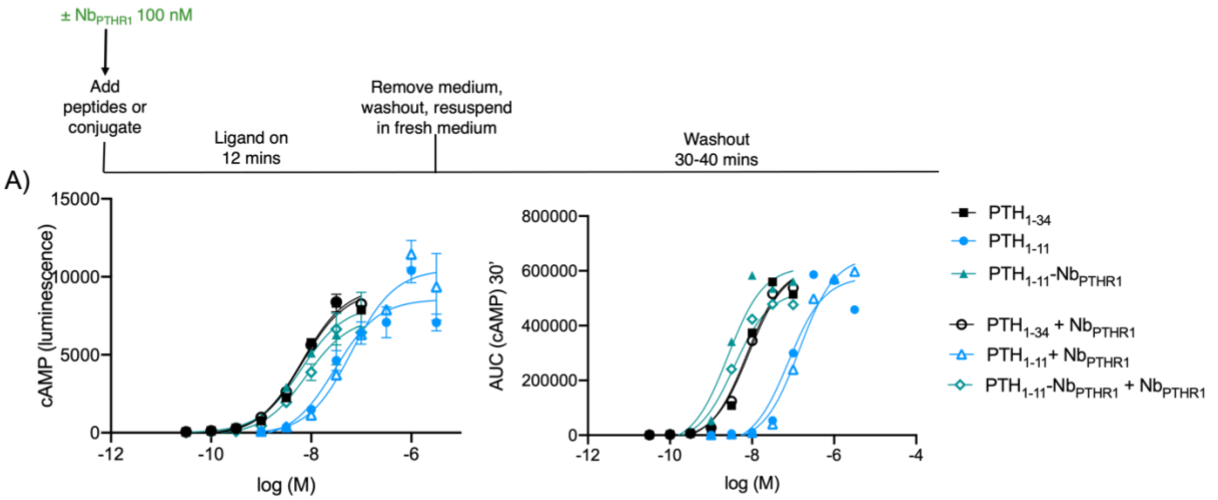


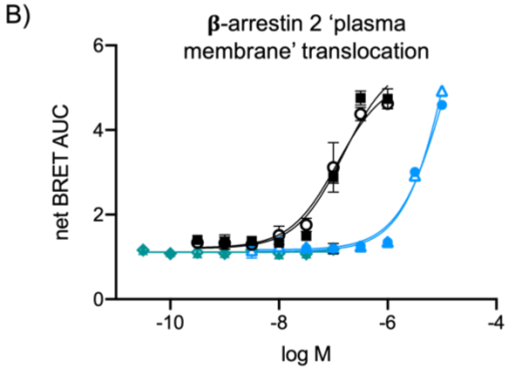


C)

| PTHR1 EC50 (±SEM) w/o $Nb_{PTHR1}$ | cAMP | Washout AUC | β-arrestin 2 (plasma membrane) |
|---|---|---|---|
| $PTH_{1-34}$ | 11 (7) | 6.7 (1.4) | 62 (44) |
| $PTH_{1-11}$ | 22 (11) | 70 (7) | 11000 (940) |
| $PTH_{1-11}$-$Nb_{PTHR1}$ | 2.8 (1) | 2.4 (0.3) | - |
| **with $Nb_{PTHR1}$** | | | |
| $PTH1_{-34}$ | 15 (11) | 5.3 (1.3) | 54 (32) |
| $PTH_{1-11}$ | 32 (16) | 115 (11) | 12000 (1090) |
| $PTH_{1-11}$-$Nb_{PTHR1}$ | 3.5 (1.4) | 3.7 (1.4) | - |

**Supporting Figure 16: Evaluation of the effects of $Nb_{PTHR1}$ co-administration on ligand signaling properties at PTHR1.** A) Timeline for the addition of ligands and Nbs in the cAMP assay. Concentration-response data for ligand-conjugates for cAMP production were assessed on WT PTHR1 with or without $Nb_{PTHR1}$ (100 nM). Data at right correspond to ligand washout responses generated from quantifying the area under the curve for kinetic signals following washout. Data points correspond to mean ± SD from technical replicates in a single representative experiment. Curves were generated using a three-parameter logistic sigmoidal model. B) Representative curve for stimulation of β-arrestin 2 recruitment to plasma membrane on WT PTHR1 with or without $Nb_{PTHR1}$. C) Tabulation of compiled agonist potency parameters for ligand-conjugates at PTHR1. $EC_{50}$ values correspond to mean (±SEM) measurements from 3 biological replicates. Note that these data are distinct from those shown in Table 1 in the main text.

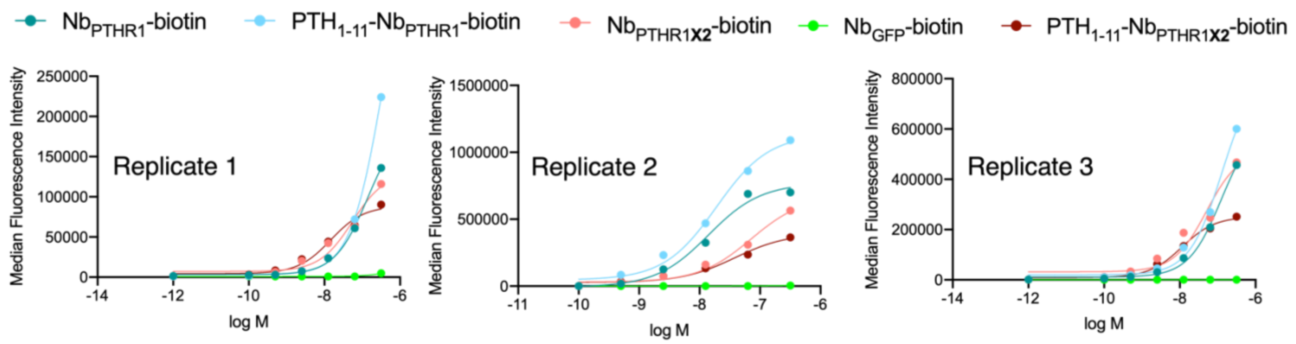


**Supporting Figure 17: Flow cytometry analysis comparing the binding of Nb and $PTH_{1-11}$-Nb conjugates to PTHR1**. Varying concentration of Nbs and ligand-conjugates labeled with biotin via sortagging were incubated with cells expressing PTHR1 followed by washing, detection with streptavidin-APC, and assessment of cellular fluorescence. $Nb_{GFP}$ refers to negative control Nb that binds to GFP not present in this receptor construct[3]. Each graphic corresponds to an independent replicate.

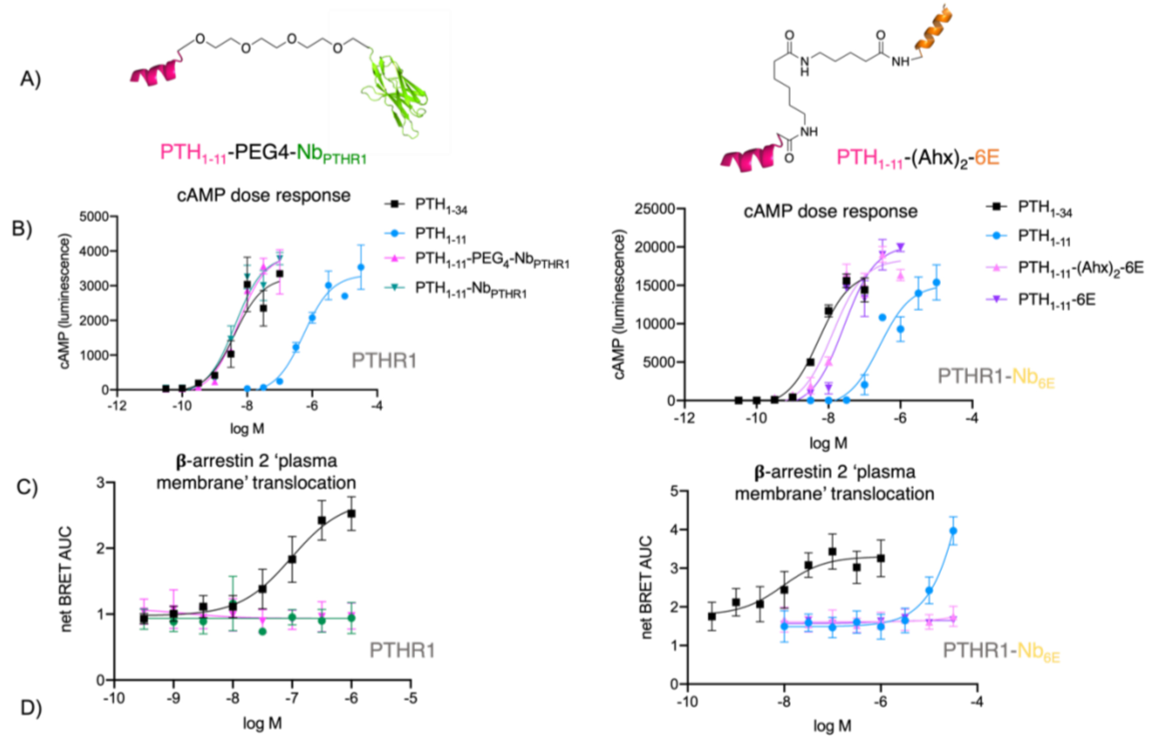


| Ligands EC$_{50}$ (±SEM) PTHR1 | cAMP | β-arrestin 2 (plasma membrane) |
|---|---|---|
| PTH$_{1-34}$ | 3.3 (3) | 87 (47) |
| PTH$_{1-11}$ | 370 (160) | ND |
| PTH$_{1-11}$-PEG$_4$-Nb$_{PTHR1}$ | 2.4 (1) | - |
| PTH$_{1-11}$-Nb$_{PTHR1}$ | 18 (15) | - |

| Ligands EC$_{50}$ (±SEM) PTHR1-Nb$_{6E}$ | cAMP | β-arrestin 2 (plasma membrane) |
|---|---|---|
| PTH$_{1-34}$ | 3.1 (1.2) | 26 (10) |
| PTH$_{1-11}$ | 140 (64) | 11000 (6500) |
| PTH$_{1-11}$-(Ahx)$_2$-6E | 9.4 (3) | - |
| PTH$_{1-11}$-6E | 12 (6) | - |

**Supporting Figure 18: Impact of longer linker length conjugates signaling activity.** A) Schematic representation of the conjugates with longer linkers used for experiments in this Figure. Compounds were synthesized as described in the methods section. Concentration-response curves for B) cAMP production, and C) β-arrestin 2 recruitment in HEK cells expressing PTHR1 (left) and PTHR1-Nb$_{6E}$ (right). D) Compiled tabulation of agonist potency parameters for conjugates with longer linkers at PTHR1 and PTHR1-Nb$_{6E}$. EC$_{50}$ values correspond to mean (±SEM) measurements from 3 independent experiments. "ND" indicates that the response was not measured in these assays. Note that these data are distinct from those in Table 1 in the main text.

>Nb$_{PTHR1}$

EVQLVESGGG LVQAGGSLRL SCAASGNIFA NNIMGWYRQP PGKEREFVAH VSHDGDSMYA

VSVKGRFAIS RKDATNLYLQ MNSLKPEDTA IYFCRLLNIP TQGRMEGFWG QGTQVTVSS

>Nb$_{PTHR1X2}$

EVQLVESGGG LVQAGGSLRL SCAASGLTFS NYAMGWFRQA PGKEREWVAS INWSGGSTYY

EDSVEGRFTI SRDNAKNTVN LQMNSLKPED TAVYYCAAKR GHYSREYDYW GQGTQVTVSS

>Nb$_{6E}$

QVQLQESGGG LVQPGGSLRL SCAASGFVFE NSAMAWYRQA PGKERELIAV IGTTFIKLAE

SVKGRFTISR DNAKSTVYLQ MNNLKPEDTA VYYCSKSGAY WGQGTQVTVS S

```
NbPTHR1       EVQLVESGGGLVQAGGSLRLSCAASGNIFANNIMGWYRQPPGKEREFVAHVSHDGD-SMY   59
NbPTHR1X2     EVQLVESGGGLVQAGGSLRLSCAASGLTFSNYAMGWFRQAPGKEREWVASINWSGGSTYY   60
Nb6E          QVQLQESGGGLVQPGGSLRLSCAASGFVFENSAMAWYRQAPGKERELIAVIGTT--FIKL   58
              :*** ******** ************  * *  *.*:** ****** :* :.

NbPTHR1       AVSVKGRFAISRKDAT-NLYLQMNSLKPEDTAIYFCRLLNIPTQGRMEGFWGQGTQVTVS 118
NbPTHR1X2     EDSVEGRFTISRDNAKNTVNLQMNSLKPEDTAVYYCAAKRG-HYSREYDYWGQGTQVTVS 119
Nb6E          AESVKGRFTISRDNAKSTVYLQMNNLKPEDTAVYYCSKS--------GAYWGQGTQVTVS 110
                **:***:***.:*. .: ****.*******:*:*             :**********

NbPTHR1       S 119
NbPTHR1X2     S 120
Nb6E          S 111
              *
```

**Supporting Figure 19: Sequence information and annotated sequence alignment data for Nbs used in this study**. Alignment was performed using ClustalOmega using the entire sequence for each Nb. Colored lettering indicates different amino acids within the sequence. "*" indicates an exact match, ":" indicates a partial match, "." indicates no match. Amino acids are numbered according to their position within the sequence.

A)

| Peptide | Sequence |
|---|---|
| PTH$_{1-28}$ | UVUEIQLMHQhRAKWLNSMRRVEWLRKKL |
| PTH$_{1-21}$ | AVUEIQLMHQhRAKWLNSMRRV |
| PTH$_{1-11}$ | ACPCVUEIQLMHQhR |
| PTHrP$_{7-36}$ | LLHDLdWKSIQDLRRRFWLHHLIAEIHTAEY |

B)

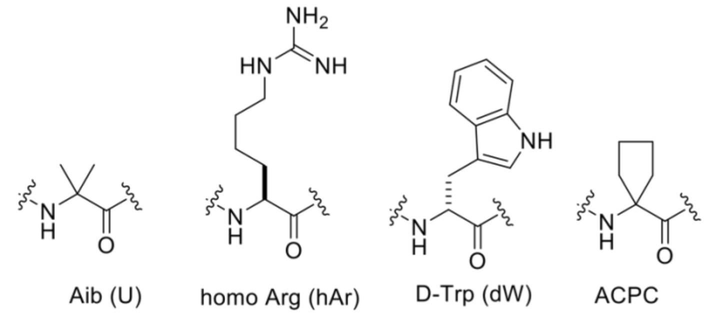


**Supporting Figure 20**: **Sequences of selected peptides used in competition binding assays.** A) Colored lettering indicates different unnatural amino acids in the peptide. B) Structures of amino acid residues presented in colored lettering.

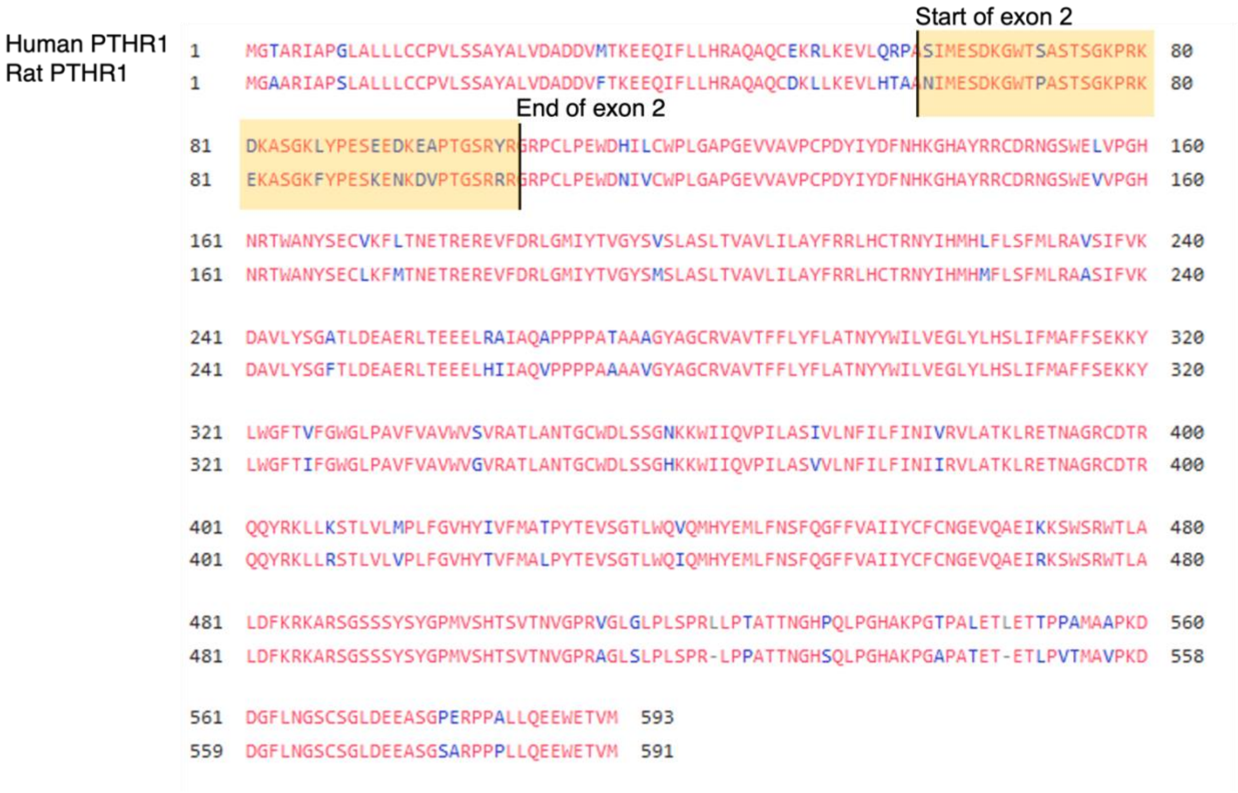


**Supporting Figure 21: Amino acid alignment of human and rat PTHR1**. The sequence from exon 2 is highlighted in yellow. Red lettering indicates conservation. Blue lettering indicates no conservation. Alignment was performed using ClustalOmega. Amino acids are numbered according to their position within the sequence.

GDDVMTKEEQIFLLHRAQAQCEKRLKEVLQRPASIMESDKGWTSASTSGKPRKDKASGKLY
PESEEDKEAPTGSRYRGRPCLPEWDHILCWPLGAPGEVVAVPCPDYIYDFNHKGHAYRRC
DRNGSWELVPGHNRTWANYSECVKFLTNETREREVFDRL*

**Supporting Figure 22: Sequence for PTHR1 extracellular domain construct used in this study.**

### Supporting Methods:

#### Assessment of receptor trafficking using fluorescence microscopy

Trafficking of PTHR1-6E was visualized as previously described[4]. Visualization was performed using a Nikon Eclipse Ni confocal microscopy. Briefly, cells expressing PTHR1-6E were seeded on glass coverslips for 24h. Staining was performed in HBSS solution supplemented with 10 mM HEPES pH 7.4 and 0.1% BSA on ice. The cells were then incubated with peptide or Nb-$PTH_{1-11}$ conjugates at 0 or 21°C for 30 min. Following washing, the cells were fixed with 4% formalin prepared in phosphate-buffered saline (PBS) and permeabilized with 0.5% Triton X-100 for 10 min. Cells were then rinsed and mounted with Everbrite Hardset Mounting Medium with DAPI (Biotium 23004).

#### Receptor internalization assay (whole cell ELISA)

Changes in the levels of cell surface receptor (PTHR1-6E) were determined in at least three independent experiments, performed in technical duplicates, using whole cell ELISA. Cells expressing PTHR1-6E (or negative control cells) were seeded into poly-L-Lysine-coated 6-well plates and incubated overnight at 37°C. Treatment compounds were prepared in cell culture medium to achieve desired concentrations. Compounds were added to cells, which were incubated for 30 min at 37°C. After incubation and washing, receptor trafficking was quenched by placing cells on ice and performing fixation with 4% paraformaldehyde in PBS for 15 min. Fixed cells were washed twice with PBS and blocked with 2% BSA in PBS for 1 h at RT. Cell surface PTHR1-6E was detected using $Nb_{6E}$-biotin (30 nM) diluted in blocking solution. $Nb_{6E}$-biotin was incubated with fixed cells for 30 min. Following washing, horseradish peroxidase-conjugated streptavidin (Pierce #21130, 1:2000 dilution) was applied to the cells and incubated for 30 min at RT. After washing, the cells were exposed to one-step TMB-ELISA solution (Thermo Scientific #34028) and incubated until visible color developed. 100 µL of developed solution was transferred to a 96-well plate, and analyzed by recording absorbance at 405 nm using a microplate reader.
